# Inhibitory input directs astrocyte morphogenesis through glial GABA_B_R

**DOI:** 10.1101/2023.03.14.532493

**Authors:** Yi-Ting Cheng, Estefania Luna-Figueroa, Junsung Woo, Hsiao-Chi Chen, Zhung-Fu Lee, Akdes Serin Harmanci, Benjamin Deneen

## Abstract

Communication between neurons and glia plays an important role in establishing and maintaining higher order brain function. Astrocytes are endowed with complex morphologies which places their peripheral processes in close proximity to neuronal synapses and directly contributes to their regulation of brain circuits. Recent studies have shown that excitatory neuronal activity promotes oligodendrocyte differentiation; whether inhibitory neurotransmission regulates astrocyte morphogenesis during development is unknown. Here we show that inhibitory neuron activity is necessary and sufficient for astrocyte morphogenesis. We found that input from inhibitory neurons functions through astrocytic GABA_B_R and that its deletion in astrocytes results in a loss of morphological complexity across a host of brain regions and disruption of circuit function. Expression of GABA_B_R in developing astrocytes is regulated in a region-specific manner by SOX9 or NFIA and deletion of these transcription factors results in region-specific defects in astrocyte morphogenesis, which is conferred by interactions with transcription factors exhibiting region-restricted patterns of expression. Together our studies identify input from inhibitory neurons and astrocytic GABA_B_R as universal regulators of morphogenesis, while further revealing a combinatorial code of region-specific transcriptional dependencies for astrocyte development that is intertwined with activity-dependent processes.

---

Astrocytes are endowed with an extraordinarily complex morphology highlighted by peripheral processes that are in close proximity to neuronal synapses^1–3^. These elaborate processes contribute to a host of synaptic functions, ultimately impacting circuit-level activities as it is estimated that a single astrocyte can interface with up to 100,000 synapses^4^. The acquisition of complex astrocyte morphologies during development is essential for the execution of these roles and has wide-ranging implications for brain function and neurological disorders^5–8^. Communication between astrocytes and neurons plays a critical role in astrocyte development^9, 10^. Previously, it was shown that structural interactions between developing astrocytes and neurons contributes to the acquisition of their complexity^11^. Moreover, astrocytes from dark-reared animals exhibit reduced territories and when coupled with evidence that glutamatergic signaling influences astrocytic volume, raise the possibility that neuronal activity itself may contribute to astrocyte morphogenesis^11–13^. Nevertheless, whether and how neuronal activity contributes to astrocyte morphogenesis remains unclear. Furthermore, what types of neurons and associated neurotransmitters provide activity-dependent input to drive astrocyte complexity are also undefined.

## Inhibitory neurons promote astrocyte morphogenesis

The acquisition of complex astrocyte morphologies in the developing cortex occurs during the P1-P28 developmental window, where the Aldh1l1-GFP reporter exhibits selective expression in developing astrocytes^14, 15^. To determine whether activation of inhibitory neurons promotes astrocyte morphogenesis, we performed intraventricular injection of AAV2/9 hDlx-hM3Dq-dTomato into P1, Aldh1l1-GFP reporter mice. One week post-injection, we treated mice with saline or 5 mg/Kg of clozapine N-oxide (CNO) two times a day, for two weeks, harvesting at P21 (**Fig. 1a**)^16^ and used slice recordings to confirm increased activity in inhibitory neurons after CNO treatment (**Fig. 1b**). We did not observe any overt differences between the saline and CNO groups with respect to astrocyte numbers (**Extended Data Fig. 1a-b**). To evaluate morphological complexity, we imaged Aldh1l1-GFP expressing astrocytes from LII-LIII of the visual cortex, finding that astrocytes from the CNO group exhibit an increase in their complexity, branch points, and process length compared to controls (**Fig. 1c-e**). CNO treatment alone had no impact on the morphological complexity of astrocytes (**Extended Data Fig. 1c**). Using the same stimulation paradigm and harvesting at P60 did not reveal any differences in astrocyte morphology, indicating that increases in complexity reflect accelerated morphogenesis (**Extended Data Fig. 1f-g**). Next, we examined whether interneuron activity is necessary for astrocyte morphogenesis by inhibiting their activity. Similar to the above studies we performed intraventricular injection of AAV2/9 hDlx-hM4Di-dTomato into P1, Aldh1l1-GFP reporter mice, treated with CNO, and harvested at P21; slice recordings confirmed decreased activity in inhibitory neurons after CNO treatment (**Fig. 1b**). Analysis of astrocyte morphology in LII-LIII of the visual cortex, revealed decreased complexity in the CNO group (**Fig. 1g-i**). Together, these observations indicate that input from inhibitory neurons contributes to astrocyte morphogenesis in the developing cortex.

**Figure 1.**
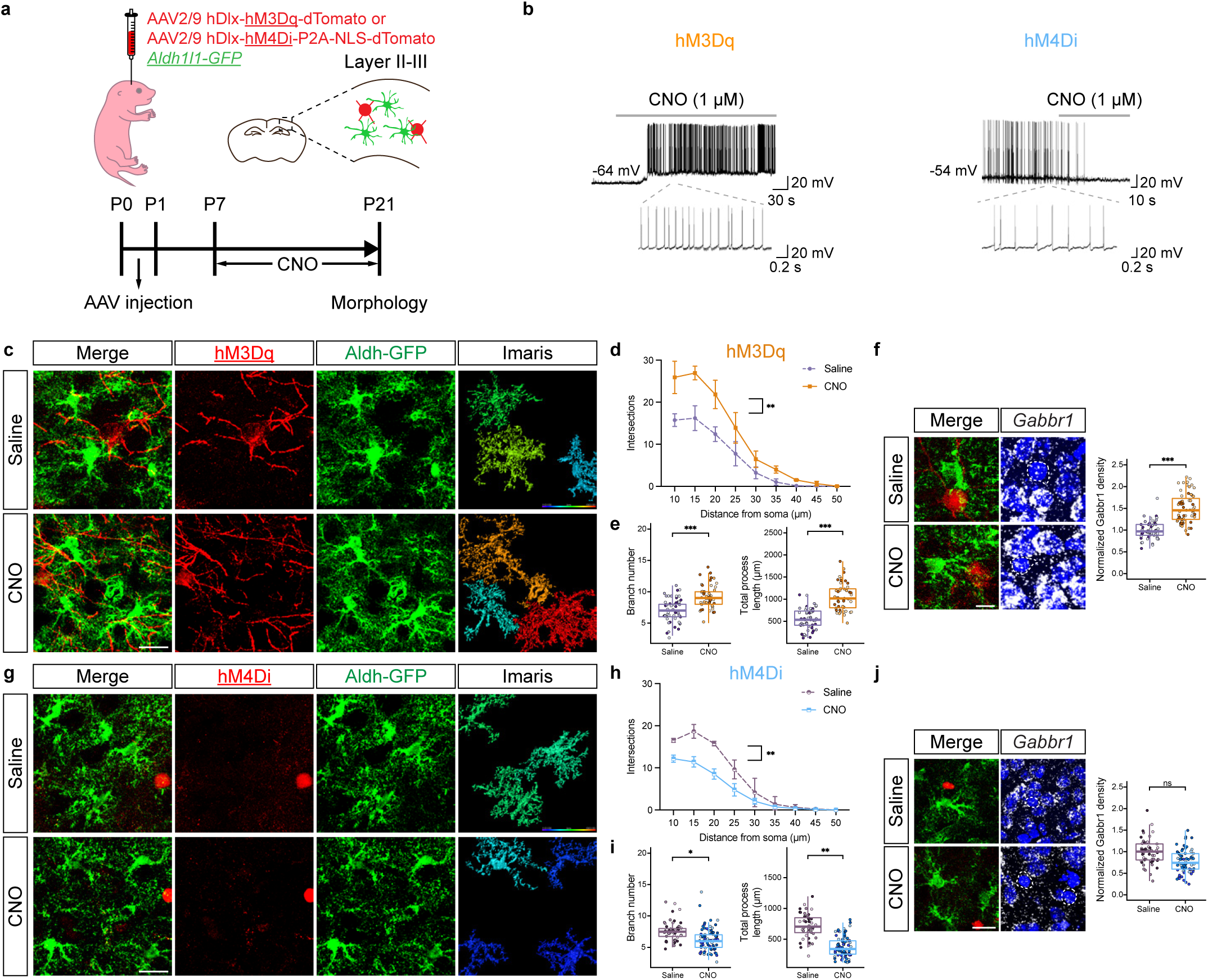
Inhibitory neuron activity regulates astrocyte morphogenesis. **a.** Schematic of DREADD-based activation of inhibitory neurons in post-natal Aldh1l1-GFP mice. **b.** Slice electrophysiological recordings of DREADD-expressing (hM3Dq or hM4Di) inhibitory neurons at P21, with and without CNO activation. Traces are representative of neuronal firing. **c-e.** Imaging of Aldh1l1-GFP astrocytes after hM3Dq activation of inhibitory neurons; quantification using Scholl analysis, branch number, and total processes at P21; *n* = 3 pairs of animals (47, 51 cells; **d**, generalized linear mixed-effects (GLME) model with Sidak’s multiple comparisons test, \*\**P* = 0.001; **e**, linear mixed-effect (LME) model, \*\*\**P* = 0.00012, 0.00013). **f.** RNAscope imaging and quantitative analysis for *Gabbr1* expression in Alhd1l1-GFP expressing astrocytes at P21; *n* = 3 pairs of animals (49, 59 cells; LME model, \*\*\**P* = 0.00090). Dashed circle denotes astrocyte with *Gabbr1*. **g-i.** Imaging of Aldh1l1-GFP astrocytes after hM4Di inhibition of inhibitory neurons; quantification using Scholl analysis, branch number, and total processes at P21; *n* = 3 and 4 animals (44, 71 cells; **h**, GLME model with Sidak’s multiple comparisons test, \*\**P* = 0.0013; **i-j**, GLME model, \**P* = 0.0327, \*\**P* = 0.0014). **j.** RNAscope imaging and quantitative analysis for *Gabbr1* expression in Alhd1l1-GFP expressing astrocytes at P21; *n* = 3 and 4 animals (49, 64 cells; LME model, *P* = 0.1014). Dashed circle denotes astrocyte with *Gabbr1*. Scale bars, 20 μm (**c**, **g**, **j**) and 10 µm (**f**). Data represent mean ± s.d. (**d**, **h**), median, minimum value, maximum value and interquartile range (IQR) (**e-f** bottom, **i-j**).

GABA is the predominant neurotransmitter released by inhibitory neurons; therefore we interrogated expression of GABA-receptors in transcriptomic data from developing Aldh1l1-GFP astrocytes. This analysis identified GABA_B_ receptor (*Gabbr1*) as upregulated in astrocytes during the P1-P14 interval in the cortex, hippocampus, and olfactory bulb (OB) (**Extended Data Figs. 1e**). To determine whether *Gabbr1* expression is associated with astrocyte morphogenesis, we performed RNAscope on P21 cortical sections from CNO and saline groups, finding that its expression is increased in Aldh1l1-GFP astrocytes when inhibitory neurons are activated (**Fig. 1f,j**). These data implicate astrocytic *Gabbr1* as a prospective regulator of astrocyte morphogenesis during development,

## *Gabbr1* regulates astrocyte morphogenesis

These observations raise the question of whether astrocytic *Gabbr1* directly regulates astrocyte morphogenesis during development. While the role of *Gabbr1* in neurons is established, whether it contributes to astrocyte development is unknown^17, 18^. To examine the role of *Gabbr1* in astrocyte development in the brain, we generated the *Gabbr1^fl/fl^; Aldh1l1-CreER; Aldh1l1-GFP* (*Gabbr1-cKO*) mouse line^19^, with the Aldh1l1-CreER line specifically targeting astrocytes (**Extended Data Fig. 2e-h**). Treatment with a single injection of tamoxifen at P1 led to efficient knockout across a host of brain regions at P28 (**Fig. 2a-b**), having no effect on the number of Aldh1l1-GFP/Sox9^+^ astrocytes at P28 in the cortex, OB, hippocampus (CA1), and brainstem (**Extended Data Fig. 3f-g**). Next, we assessed morphological complexity of astrocytes from the *Gabbr1-cKO* at P28, focusing on layer II-III (LII-LIII) of the visual cortex, external plexiform layer (EPL) OB, CA1 in the hippocampus, the internal granule layer (IGL) in the cerebellum, and the medulla in the brainstem. We found that knockout of *Gabbr1* led to a reduction in astrocyte complexity, branch points, and process length in all the examined brain regions (**Fig. 2c and Extended Data Fig. 3a-b**); these observations were validated using sparse labeling and knockout of *Gabbr1* (**Extended Data Fig. 3c-e**). Together, these data indicate that *Gabbr1* is a universal regulator of astrocyte morphogenesis. Next, we evaluated spontaneous Ca2^+^ activity in the cortex of *Gabbr1-cKO* and control mice at P28. Using a floxed-dependent GCaMP6 mouse line within our *Gabbr1-cKO* line (**Fig. 2a**), followed by *ex vivo*, two-photon slice imaging at P28, we found no changes in spontaneous Ca2^+^ activity in cortical astrocytes from the *Gabbr1-cKO* (**Fig. 2d; Extended Data Fig. 6a**), suggesting physiological activities of astrocytes are unaffected. Upon binding to GABA, astrocytic *Gabbr1* elicits Ca2^+^ responses^20^, therefore we treated slices with baclofen, the GABA_B_ receptor agonist, and found that cortical astrocytes from the *Gabbr1-cKO* failed to generate a baclofen-induced Ca2^+^ response (**Fig. 2e**). These data suggest that inhibitory input is disrupted in *Gabbr1-cKO* astrocytes and in conjunction with our cellular analysis indicate that astrocytic *Gabbr1* regulates morphogenesis.

**Figure 2.**
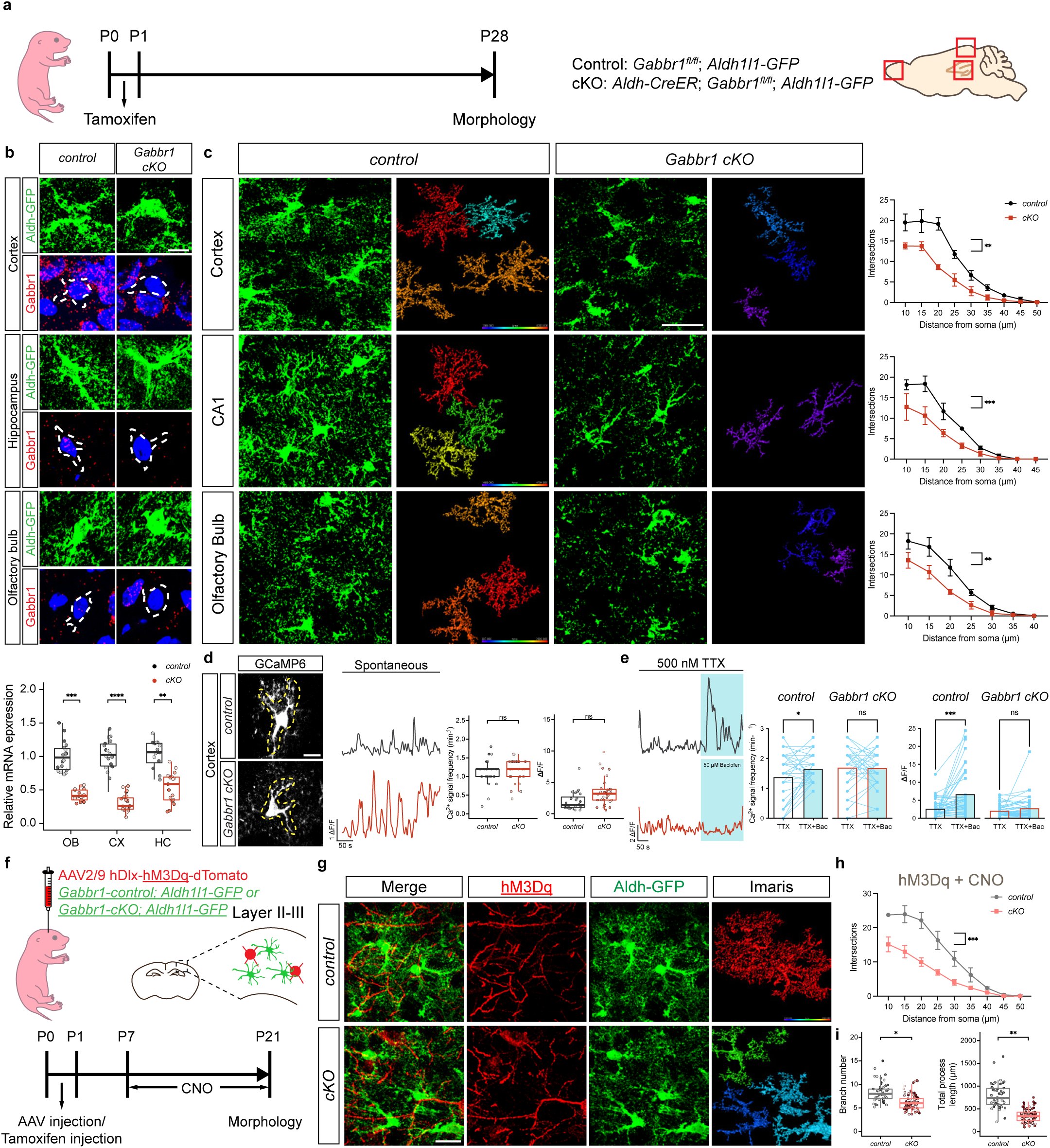
*Gabbr1* is required for astrocyte morphogenesis. **a.** Experimental timeline and mouse lines rendering astrocyte-specific knockout of *Gabbr1*. **b.** RNA-Scope imaging of *Gabbr1* within Aldh1l1-GFP astrocytes from control and *Gabbr1-cKO* mouse lines; quantification derived from *n* = 3 pairs of animals (*control*: OB 18, CX 18, HC 16; *cKO*: OB 17, CX 19, HC 17 cells; LME model, \*\*\**P* = 0.00020, \*\*\*\**P* = 3.07e-06, \*\**P* = 0.0030). Dashed circle denotes astrocyte with *Gabbr1*. **c.** Imaging of Aldh1l1-GFP astrocytes from the cortex, CA1 of the hippocampus, and olfactory bulb at P28; quantification via Scholl analysis derived from *n* = 3 pairs of animals (*control*: OB 43, CX 27, HC 32, *cKO*: OB 31, CX 33, HC 30 cells; GLME model with Sidak’s multiple comparisons test, \*\**P* = 0.0011, *** *P* = 0.0006, \*\**P* = 0.0037). **d.** Imaging of GCaMP6s activity in *control* and *Gabbr1-cKO* astrocytes from the cortex at P28; quantification is derived from *n* = 3 pairs of animals (24,33 cells; GLME model, *P* = 0.6361, 0.2239). **e.** Imaging of GCaMP6s activity in the presence of TTX and baclofen; quantification derived from *n* = 40 cells from 3 pairs of animals (two-tailed Wilcoxon matched-pairs signed rank test, \**P* = 0.022, *P* = 0.89, \*\*\**P* = 0.0006, *P* = 0.32). **f.** DREADD-based activation of inhibitory neurons in *Gabbr1-cKO* mice. **g-i.** Imaging of Aldh1l1-GFP astrocytes from *Gabbr1-cKO* (and *control*) after hM3Dq activation of inhibitory neurons; quantification using Scholl analysis, branch number, and total processes at P21; *n* = 3 and 5 animals (50, 80 cells; **h**, GLME model with Sidak’s multiple comparisons test, \*\**P* = 0.0011, **i**; GLME model, \**P* = 0.034, \*\**P* = 0.0026). Scale bars, 10 μm (**b, d**), 30 µm (**c**), and 20 μm (**g**). Data represent mean ± s.d. (**c**, **h**), median, minimum value, maximum value and IQR (**b**, **d**, **i**).

Next, we examined whether inhibitory input functions through *Gabbr1* to regulate astrocyte morphogenesis. To test this we injected the *Gabbr1-cKO* mouse line (and control) at P1 with AAV2/9 hDlx-hM3Dq-dTomato and treated with CNO (**Fig. 2f**). Assessing astrocyte morphogenesis at P21 revealed that activation of inhibitory neurons did not promote astrocyte morphogenesis in the *Gabbr1-cKO* (**Fig. 2g-i**) and that the extent of astrocyte complexity was similar to the *Gabbr1-cKO* (**Fig. 2c v 2g-i**). Collectively, these data indicate that inhibitory neurons drive astrocyte morphogenesis through astrocytic *Gabbr1*.

## Astrocytic *Gabbr1* regulates cortical circuits

The defects in astrocyte morphogenesis in the *Gabbr1-cKO* prompted us to perform single-cell RNA Sequencing (scRNA-Seq) on *Gabbr1-cKO* and control cortices from P28. Using Seurat analysis we identified the principle cell types in the brain and did not observe any difference in their constituency (**Extended Data Fig. 4a-b**)^21^. Next, we used the CellChat pipeline to map cell-cell interactions between astrocytes and excitatory- and inhibitory-neurons from the scRNA-Seq datasets^22^. This analysis revealed a decrease in the number of interactions between astrocytes and excitatory neurons, coupled with an increase in the interaction between astrocytes and inhibitory neurons in the *Gabbr1-cKO* cortex **(Fig. 3a; Extended Data Fig. 4c-e**). KEGG pathway analysis of the differentially expressed genes (DEGs) in neurons revealed dysregulated expression of GABAergic synapses, suggesting alterations in astrocyte-neuron communication in the *Gabbr1-cKO* cortex **(Fig. 3b-c; Extended Table 3**).

**Figure 3.**
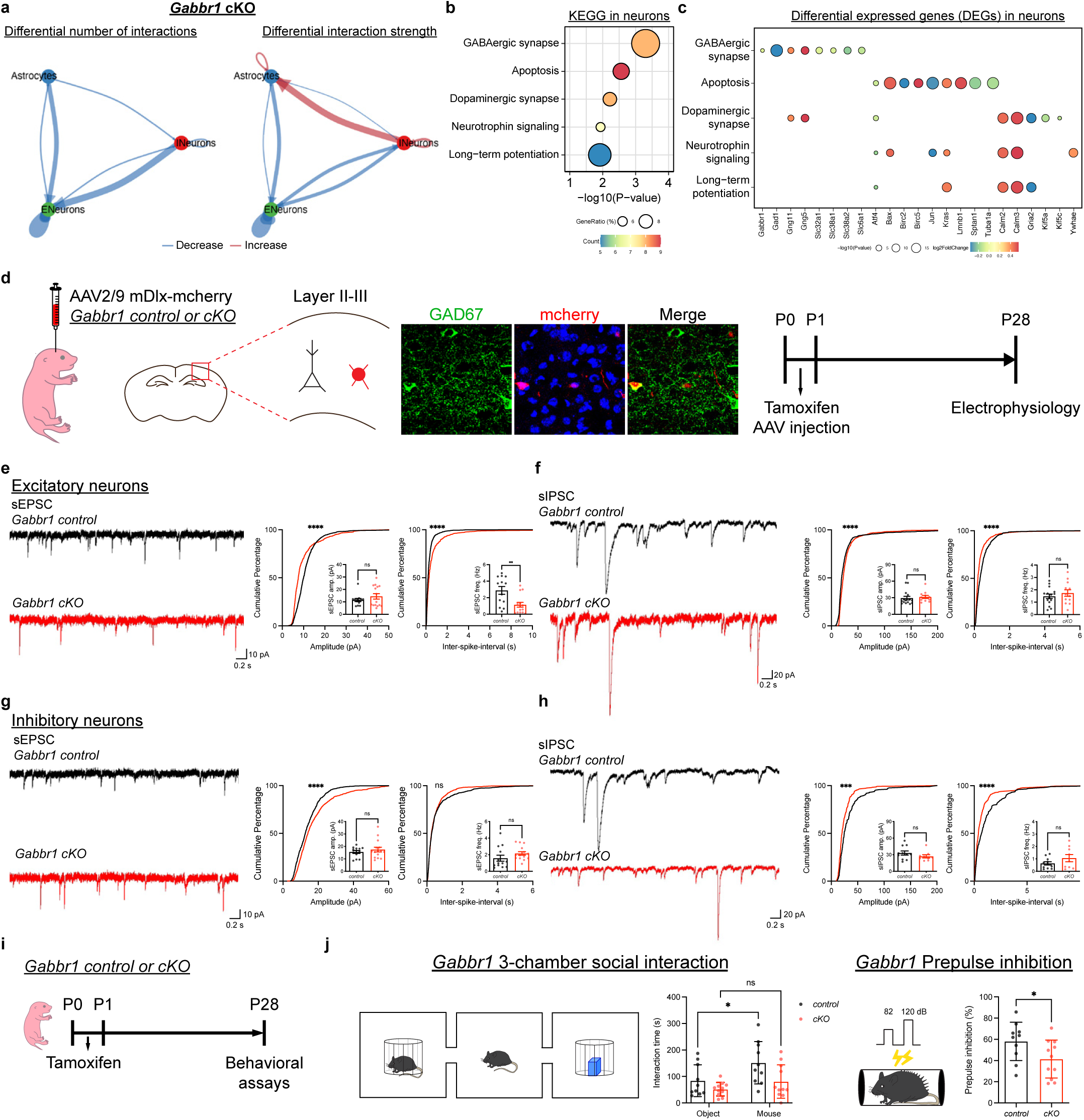
Loss of astrocytic *Gabbr1* disrupts cortical circuit function. **a.** CellChat interaction diagram illustrating astrocyte interactions with neurons in the cortex from P28 *Gabbr1-cKO* mice; width of colored arrow indicates scale of interaction. See Extended Data Figure 4. **b-c**. KEGG pathway analysis of neurons from *Gabbr1-cKO* scRNA-Seq (**b**, analyzed by Enrichr) and dot plot of differentially expressed genes from KEGG (**c**, analyzed by Seurat FindMarkers). **d.** Schematic of viral labeling of inhibitory neurons and experimental timeline. **e-h.** Representative traces of spontaneous EPSCs and IPSCs from excitatory and inhibitory neurons from cortex of *Gabbr1-cKO* and *controls*. Associated cumulative and bar plots demonstrate quantification of sEPSC and sIPSC from 3 pairs of animals (**e**, *n* = 13, 15 cells; Kolmogorov-Smirnov test, **** *P* < 0.0001; two-tailed Mann-Whitney test, *P* = 0.8207, \*\**P* = 0.003; **f**, *n* = 15, 12 cells; Kolmogorov-Smirnov test, **** *P* < 0.0001; two-tailed Mann-Whitney test, *P* = 0.1995, 0.5888; **g**, *n* = 13, 15 cells; Kolmogorov-Smirnov test, **** *P* < 0.0001; two-tailed Mann-Whitney test, *P* = 0.7856, 0.0504; **h**, *n* = 11, 9 cells; Kolmogorov-Smirnov test, **** *P* < 0.0001; two-tailed Mann-Whitney test, *P* = 0.2299, 0.3796). **i.** Experimental timeline for behavioral analysis. **j**. 3-chamber social interaction and pre-pulse inhibition studies on *Gabbr1-cKO* and *control* mice from 10 animals in control group and 11 animals in cKO group (left, GLME model with Sidak’s multiple comparisons test, \**P* = 0.015; right, two-tailed Mann-Whitney test, \**P* = 0.043). Data represent mean ± s.e.m. (**e-h**), s.d. (**j**).

To validate these findings, we quantified excitatory and inhibitory synapses in LI and LII-II, respectively, from *Gabbr1-cKO* mice at P28. This analysis revealed an increase in excitatory Vglut2/Psd95 synapses, coupled with no changes in the number of inhibitory vGat/Gephrin synapses (**Extended Data Fig.5a-g)**. These changes in synaptic numbers led us to evaluate whether loss of astrocytic *Gabbr1* influences neuronal activity. Using intraventricular injection of AAV2/9-mDlx-mRuby2 at P1 to label interneurons in *Gabbr1-cKO* and control mice (**Fig. 3d**), we evaluated neuronal excitability, finding no differences in action potential firing between cKO and control groups (**Extended Data Fig. 6b-i**). Next, we measured synaptic transmission through spontaneous excitatory postsynaptic current/inhibitory postsynaptic current (sEPSC/IPSC) finding dysregulation of both excitatory and inhibitory activity in LII-LIII neurons. Excitatory neurons exhibited decreased sEPSC activity via cell average and K-S test, while exhibiting no significant difference in sIPSC activities in cell averages and a significant difference via K-S test (**Fig. 3e-f**). Analysis of inhibitory neurons revealed increased sEPSC amplitudes and decreased sIPSC amplitudes via K-S test, which were not statistically significant when averaged across cells (**Fig. 3g-h**). Next, we subjected the *Gabbr1-cKO* (and control) mice to a series of behavioral tests, identifying deficits in pre-pulse inhibition and three-chamber social interaction in the *Gabbr1-cKO* mice (**Fig. 3i-j**; **Extended Data Fig.5h-m**). Collectively, these molecular, physiological, and behavioral data indicate that astrocytic *Gabbr1* mediates interactions with excitatory and inhibitory neurons that contributes to functioning cortical circuits.

## *Gabbr1* regulates *Ednrb1* during astrocyte morphogenesis

To identify the mechanisms downstream of *Gabbr1* regulating astrocyte morphogenesis we performed bulk RNA-Seq on FACS purified astrocytes from P28 *Gabbr1-cKO* mice from the cortex, hippocampus, olfactory bulb (**Fig.4a; Extended Tables 4**). Gene Ontology (GO) analysis of the DEGs in *cKO* astrocytes highlighted extra-cellular matrix and membrane-associated genes as the most represented across these regions (**Fig. 4b-c**). From this group, we focused on Endothelial Receptor B (*Ednrb*) and confirmed reduced expression in astrocytes from the *Gabbr1-cKO* mouse (**Fig. 4d-e**). *Ednrb* is a GPCR that regulates cytoskeletal dynamics through Ca^2+^ activity and actin organization in astrocytes^23, 24^ and contributes to reactive astrocyte responses after brain injury^25^, however its role in astrocyte morphogenesis is unknown. To examine whether *Ednrb* regulates astrocyte morphogenesis, we employed the *Rosa-LSL-Cas9-eGFP* mouse line, along with AAV-approaches to express Cas9 in astrocytes and guideRNAs targeting *Ednrb* (**Fig. 4f**), which enabled selective deletion of *Ednrb* in cortical astrocytes (**Fig.4g-i and Extended Data Fig. 10b,f).** Using the mCherry tag on the AAV-GFAP-Cre virus to assess morphology in astrocytes that had lost *Ednrb*, we found a reduction in morphological complexity (**Fig. 4g-i**). These findings highlight a new role for *Ednrb* in astrocyte morphogenesis in the cortex and identify molecular processes that act downstream of *Garbbr1*.

**Figure 4.**
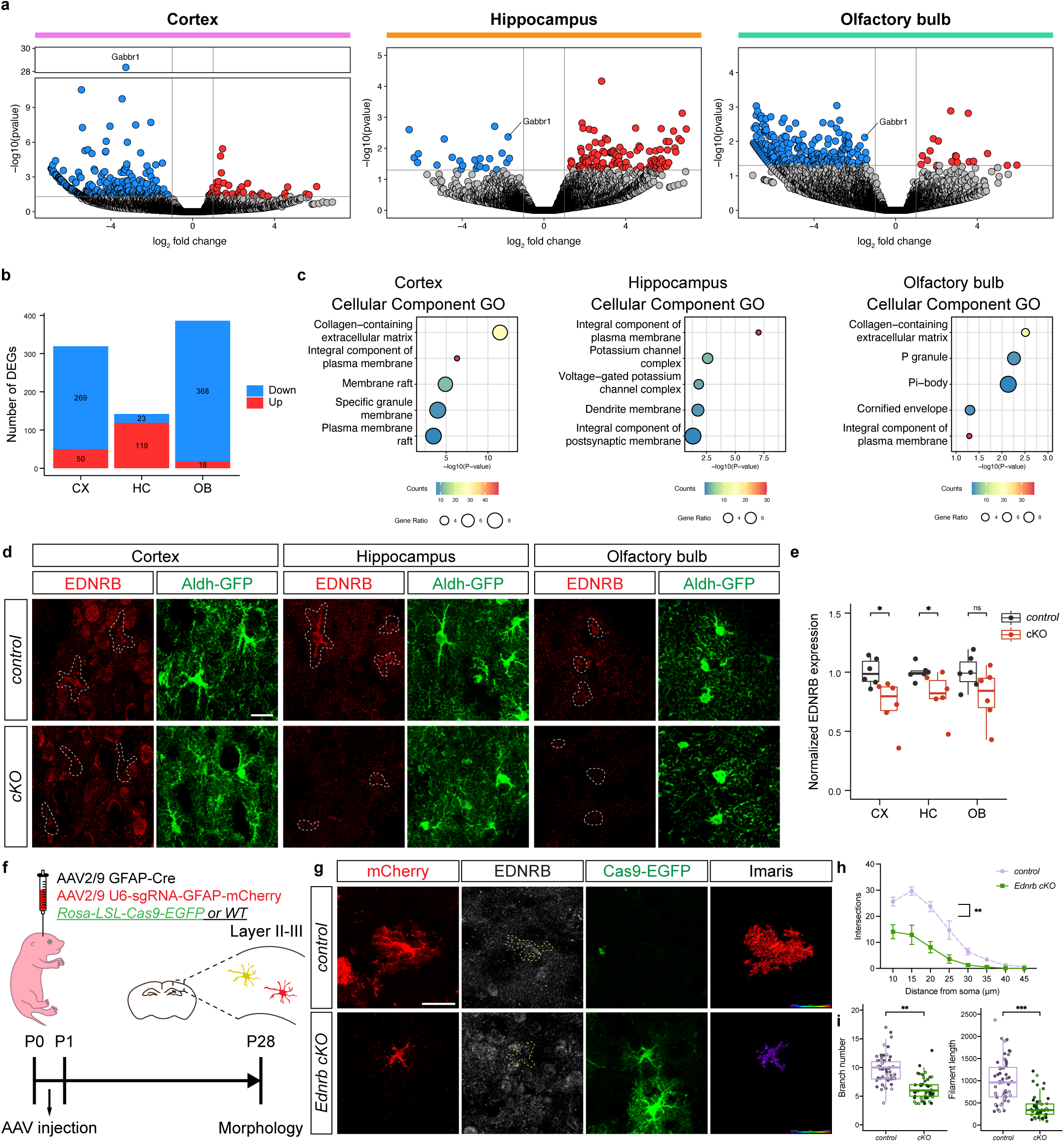
*Gabbr1* regulates astrocyte morphology through *Ednrb1*. **a.** Volcano plots from RNA-Seq analysis of control and *Gabb1-cKO* astrocytes from cortex, hippocampus, and OB. **b.** Table of the number of differentially expressed genes (DEGs) from each region. **c.** Gene Ontology (GO) analysis of DEGs performed with Enrichr. **d.** Immunostaining for EDNRB in P28 astrocytes from *Gabbr1-cKO* and *control* astrocytes. **e.** Quantification of EDNRB expression in *Gabbr1-cKO* and control from *n* = 6 pairs of animals (two-tailed Mann-Whitney test, \**P* = 0.041, 0.015, *P* = 0.1320). **f.** Schematic and timeline of selective deletion of *Ednrb* in cortical astrocytes. **g-i.** Imaging of virally labeled astrocytes from the P28 cortex of mice where *Ednrb* has been knocked out using guideRNAs in the *ROSA-LSL-Cas9-EGFP* mouse line; quantification via Scholl analysis was derived from *n* = 3 pairs of animals (53, 49 cells; **h**, GLME model with Sidak’s multiple comparisons test, \*\**P* = 0.001; i, GLME model, \*\**P* = 0.001, \*\*\**P* = 0.0002). Scale bars, 20 μm (**d**), 30 μm (**g**). Data represent mean ± s.d. (**h**), median, minimum value, maximum value and IQR (**e**, **i**).

## Region-specific regulation of *Gabbr1*

To understand how *Gabbr1* fits into astrocytic developmental programs, we sought to define the transcriptional mechanisms that control its expression. Our astrocyte transcriptomic dataset from P1-P14 in the developing brain revealed temporal and region-specific differences in gene expression profiles between the cortex, hippocampus, and OB (**Extended Data Fig. 2a-d; Extended Table 1**), suggesting region-specific mechanisms may regulate Gabbr1 expression. This prompted us to perform Homer motif analysis on the DEGs between P1 and P14, identifying numerous transcription factors (TFs) whose motifs are enriched from each region (**Fig. 5a**). Next, we filtered these candidate TFs based on their expression levels, which nominated *Nfia and Sox2* in the cortex, *Sox9* and *Nr2f1* in the hippocampus, and *Sox9* and *Tead1* in the OB (**Fig. 5a, Extended Data Fig. 2a-c**).

**Figure 5.**
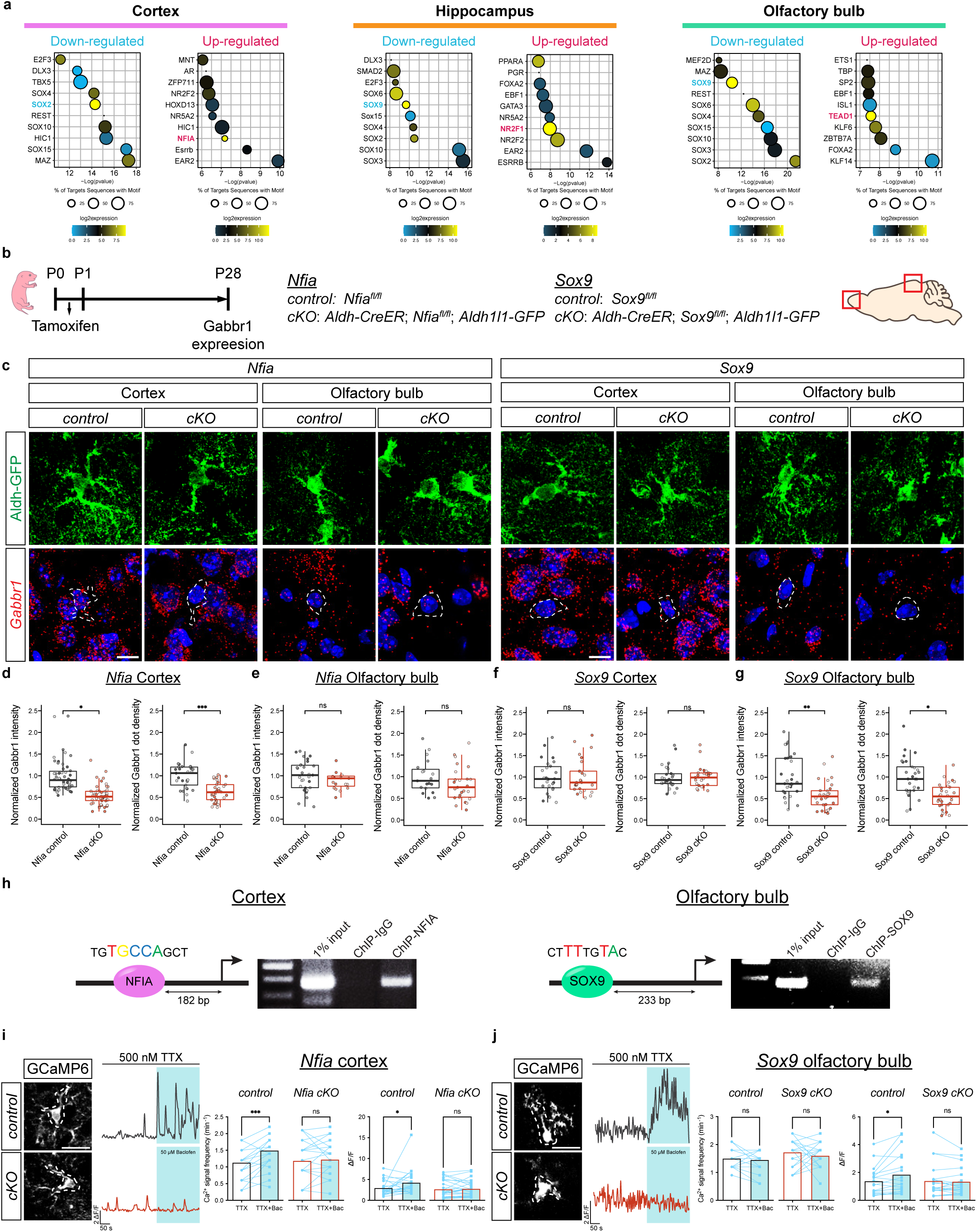
Region-specific regulation of *Gabbr1* by SOX9 and NFIA. **a.** Homer transcription factor motif analysis on differentially expressed genes (DEGs) from P1 and P14 timepoints from astrocytes isolated from the cortex, hippocampus, and olfactory bulb. **b.** Schematic depicting mouse lines and experimental timelines. **c-h.** RNAscope imaging of *Gabbr1* expression in Aldh1l1-GFP astrocytes from *Nfia-cKO*, *Sox9-cKO* and associated controls at P28; quantitative analysis of *Gabbr1* expression is derived from *n* = 3 pairs of animals (**d**, 50, 50, 29, and 33 cells; GLME model, \**P* = 0.022; LME model \*\*\**P* = 0.0002; **e**, 29, 19, 19, and 25 cells; LME model, *P* = 0.17, 0.27; **f**, 26, 25, 26, and 29 cells; GLME model, *P* = 0.91, 0.95; **g**, 30, 29, 29, 30 cells; LME model, \*\**P* = 0.0094, \**P* = 0.014). Dashed circle denotes astrocyte with *Gabbr1*. **h.** Chromatin immunoprecipitation of NFIA from P28 cortex or SOX9 from P28 olfactory bulb (OB), followed by PCR detection of association with motif in proximal promoter region of *Gabbr1*. **i-j.** Imaging of GCaMP6s activity from the cortex of *Nfia-cKO* mice or the OB from *Sox9-cKO* mice (and *controls*) in the presence of TTX and baclofen; quantification is derived from *n* = 19-26 cells from 3 pairs of animals (**i**, 23, 26 cells, two-tailed Wilcoxon matched-pairs signed rank test, \*\*\**P* = 0.0001, *P* = 0.53, \**P* = 0.048, *P* = 0.37; **j**, 19 20 cells, two-tailed Wilcoxon matched-pairs signed rank test, P = 0.65, 0.45, \**P* = 0.012, *P* = 0.81). Scale bars, 10 μm (**c**) and 20 µm (**i-j**). Data represent median, minimum value, maximum value and IQR (**d-g**).

To determine whether SOX9 and NFIA regulate *Gabbr1* in developing astrocytes we utilized *Nfia^fl/fl^; Aldh1l1-CreER; Aldh1l1-GFP* (*NFIA-cKO*) and *Sox9^fl/fl^; Aldh1l1-CreER; Aldh1l1-GFP* (*Sox9-cKO*) mouse lines that enable temporal control of deletion in astrocytes^26, 27^. To delete *Sox9* or *Nfia* during astrocyte morphogenesis, we treated the above mouse lines (and *Nfia^fl/fl^*; *Aldh1l1-GFP* or *Sox9^fl/fl^*; *Aldh1l1-GFP* controls) with a single injection of tamoxifen at P1 (**Fig. 5b**); analysis at P7 and P28 revealed efficient knockout (**Extended Data Fig. 7a-d**). RNAscope analysis of *NFIA-cKO* mice revealed that *Gabbr1* is specifically downregulated in Aldh1l1-GFP astrocytes from the cortex, but not the OB (**Fig. 5c-e**). Conversely, in the *Sox9-cKO*, we found that *Gabbr1* is downregulated in Aldh1l1-GFP astrocytes from the OB and not the cortex (**Fig. 5c, f-g**). Next, we examined whether NFIA and SOX9 are sufficient to induce *Gabbr1* expression, finding that NFIA overexpression in the cortex resulted in increased *Gabbr1* expression, while SOX9 promotion of *Gabbr1* expression in the OB was not significant (**Extended Data Fig. 7g-h**). To determine if *Gabbr1* is a direct target of NFIA and SOX9, we performed chromatin immunoprecipitation PCR (ChIP-PCR) for the NFIA or SOX9 binding motifs from P28 cortex and olfactory bulb, respectively (**Fig. 5h**). These ChIP-PCR assays revealed that NFIA and SOX9 bind to their sites in the *Gabbr1* promoter in the cortex and OB. Together, these data indicate region-specific regulation of *Gabbr1* by NFIA and SOX9 in the cortex and OB, respectively.

To test whether GABA-induced Ca2^+^ responses are impaired in the cortex of the *Nfia-cKO* or OB of the *Sox9-cKO*, we used GCaMP6s and measured Ca2^+^ activity in astrocytes using ex-vivo, two photon imaging. Application of baclofen, the GABA_B_ receptor agonist, revealed that cortical astrocytes from the *Nfia-cKO* and OB astrocytes from the *Sox9-cKO* failed to generate a baclofen-induced Ca2^+^ response (**Fig. 5i-j**; **Extended Data Fig. 7e-f**). These observations indicate that *Gabbr1* responses are impaired in *Nfia* and *Sox9* mutant astrocytes from the cortex and OB, respectively, further region-specific regulation of *Gabbr1* expression.

## Region-specific regulation of astrocyte morphogenesis

*Sox9* and *Nfia* play an important role in early glial specification in the embryonic spinal cord, however whether they regulate astrocyte morphogenesis in the brain is unknown^28–30^. Recent studies have also shown that despite universal expression in astrocytes, *Sox9* is required to maintain astrocyte complexity in the adult olfactory bulb, while *Nfia* is required to maintain astrocyte complexity in the adult hippocampus and adult cortex^26, 27^. However, whether these region-specific transcriptional dependencies in the adult are developmentally encoded remains unknown.

To determine whether *Sox9* and *Nfia* regulate astrocyte morphogenesis in a region-specific manner we harvested *Sox9-cKO* and *Nfia-cKO* (and controls) at P28. Our initial analysis found no changes in proliferation or gross number of Aldh1l1-GFP astrocytes at P28 in the cortex and OB in both the *Nfia-cKO* and *Sox9-cKO* mice, respectively (**Extended Data Fig. 8e-h**). To evaluate the morphological complexity of astrocytes from the *Nfia-cKO* and *Sox9-cKO* we focused on layer II-III (LII-LIII) of the visual cortex, external plexiform layer (EPL) OB, and CA1 in the hippocampus. We found that knockout of *Sox9* led to a reduction in astrocyte complexity in the OB, whereas astrocytes in the cortex or hippocampus are unaffected (**Fig. 6a-b, Extended Data Fig. 8b,d**). In contrast, knockout of *Nfia* led to a reduction in astrocyte complexity in the hippocampus and cortex, but not the OB (**Fig. 6a-b, Extended Data Fig. 8a,c**). These data indicate that region-specific transcriptional dependencies regulate astrocyte morphogenesis during development.

**Figure 6.**
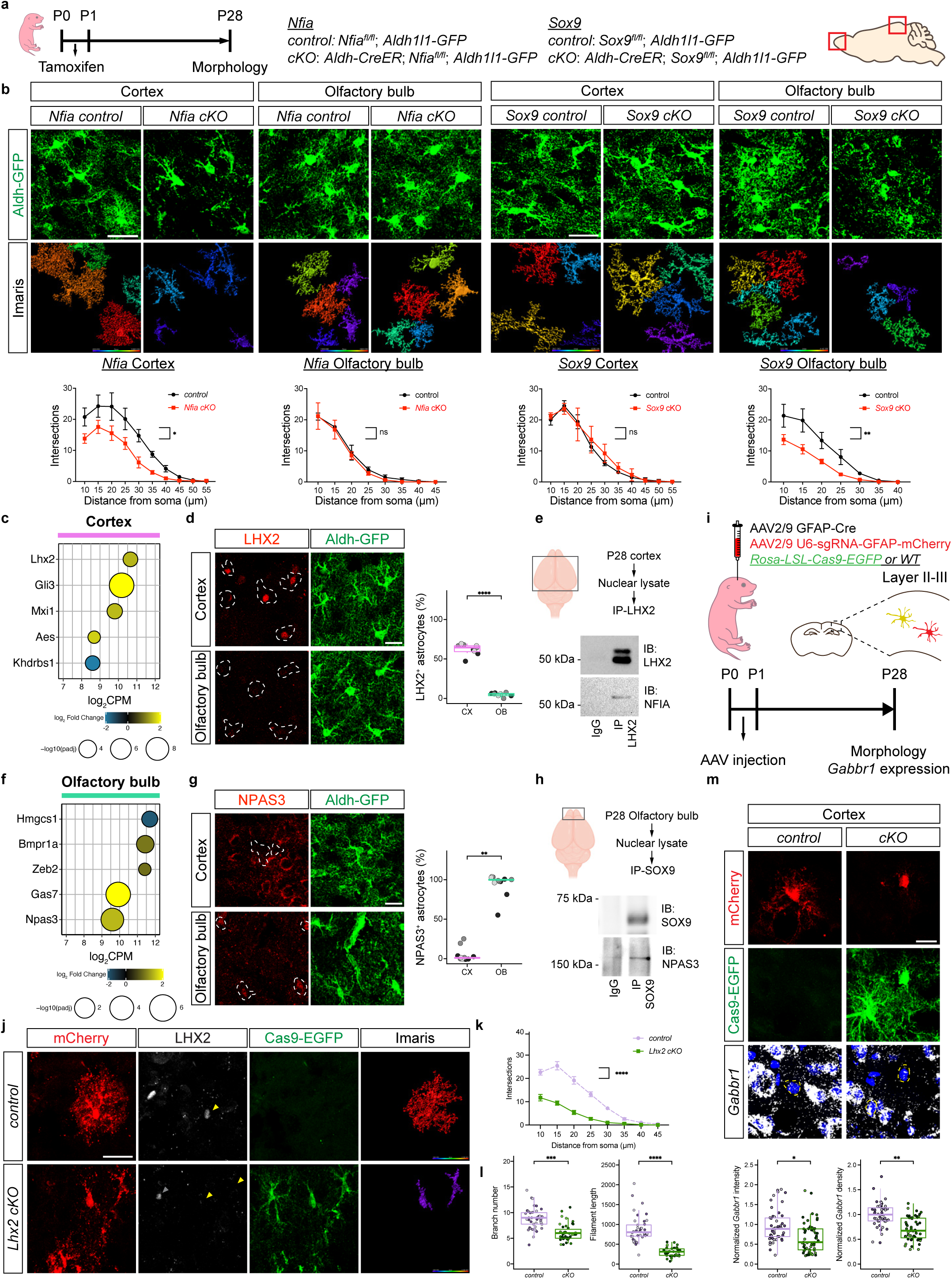
Regulation of astrocyte morphogenesis by region-specific mechanisms. **a.** Timeline and mouse lines rendering astrocyte-specific knockout of *Sox9* and *Nfia*. **b.** Imaging of Aldh1l1-GFP astrocytes at P28 from the *Sox9-cKO* and *Nfia-cKO*; quantification via Scholl analysis was derived from *n* = 3 pairs of animals (*Nfia control*: OB 56, CX 52, *Nfia cKO*: OB 60, CX 48, *Sox9 control*: OB 29, CX 35, *Sox9 cKO*: OB 39, CX 33 cells; GLME model with Sidak’s multiple comparisons test, \**P* = 0.015, *P* = 0.41, 0.60, \*\**P* = 0.0097). **c.** CX-specific DEGs increased between P7-P14. **d**. Immunostaining for LHX2 in P28 astrocytes quantified from *n* = 3 pairs of animals (LME model, \*\*\*\**P* = 1.31e-12). **e**. Immunoprecipitation of LHX2 and immunoblot of LHX2 and NFIA from the P28 cortex. **f.** OB-specific DEGs increased between P7-P14. **g**. Immunostaining for NPAS3 in P28 astrocytes quantified from *n* = 3 pairs of animals (GLME model, \*\**P* = 0.0013). **h**. Immunoprecipitation of NPAS3 and immunoblot of NPAS3 and SOX9 from P28 cortex. **i.** Schematic of *Lhx2* deletion in cortical astrocytes. **j-l**. Imaging of virally labeled astrocytes lacking *Lhx2* from P28 cortex; quantification via Scholl analysis was derived from *n* = 3 pairs of animals (41,41 cells; **k**, GLME model with Sidak’s multiple comparisons test, \*\*\*\**P* = 1.29e-24; **l**, GLME model, \*\*\**P* = 0.00099, \*\*\*\**P* = 3.98e-05). **m**. RNAscope for Gabbr1 expression in *Cas9-EGFP* cortical astrocytes lacking *Lhx2* and controls at P28; quantitative analysis demonstrating reduction of *Gabbr1* expression is derived from *n* = 3 pairs of animals (51,55 cells; GLME model, \**P* = 0.015; LME model, **P = 0.0003). Scale bars, 30 μm (**b, j**), 20 μm (**d**, **g**, **m**). Data represent mean ± s.d. (**b**, **k**), median, minimum value, maximum value and IQR (**d**, **g**, **l**, **m**).

These studies highlight a possible role for *Nfia* in the development and function of cortical circuits. To interrogate synapse formation we quantified excitatory and inhibitory synapses in LI and LII-II, finding no changes in the number of vGlut2/Psd95, vGat/Gephyrin, or vGlut1/Psd95 puncta from *NFIA-cKO* mice (**Extended Data Fig. 9a-d**). Measuring synaptic transmission through spontaneous excitatory postsynaptic current/inhibitory postsynaptic current (sEPSC/IPSC), we found decreases in sEPSC/IPSC in both excitatory and inhibitory neurons in LII-III via K-S test that were not statistically significant when averaged across cells; sIPSC of inhibitory neurons demonstrated significant decreases via K-S test and across cell averages (**Extended Data Fig. 9e-f**). Next, we subjected these mice to a series of behavioral assays finding specific defects in pre-pulse inhibition and three-chamber social interaction (**Extended Data Fig. 9g-h;k-p**), deficits that parallel our observations in the *Gabbr1-cKO* mouse line (**Fig. 3j**). Collectively, these data indicate that astrocytic NFIA contributes to the development of cortical circuits and implicates astrocyte morphogenesis as a central component of circuit maturation.

## LHX2 cooperates with NFIA to regulate cortical astrocyte morphogenesis

Because NFIA and SOX9 exhibit universal expression in astrocytes and have region-specific roles, led us to examine how this regional specialization is conferred. Identification of region-specific transcriptional mechanisms may reveal insights into the regional regulation of astrocyte morphogenesis and *Gabbr1* expression during development. Analysis of regional and temporal signatures from our developing astrocyte RNA-Seq data (**Extended Data Fig. 2a-c**) revealed a cohort of TFs expressed in cortical or olfactory bulb astrocytes (**Figs.6c,f, Extended data table 5**). We found that the transcription factor *Lhx2* is expressed in cortical astrocytes, while the transcription factor *Npas3* is expressed in olfactory bulb astrocytes (**Fig. 6d,g**). We previously demonstrated that hippocampal-specific functions of NFIA in the adult are mediated by interactions with other transcription factors^26^, therefore we examined whether LHX2 and NPAS3 interact with NFIA or SOX9, respectively. Towards this we performed a series of co-immunoprecipitation experiments, finding that NFIA associates with LHX2 in the cortex, while NPAS3 associates with SOX9 in the olfactory bulb (**Fig. 6e,h**).

Prior studies on *Lhx2* suggest that it has a region-specific role in the embryonic brain, where it promotes neurogenesis in the hippocampus by antagonizing NFIA function^31^. Interestingly, *Lhx2* does not promote neurogenesis in the cortex and its role in astrocyte development remains unknown. Using the *Rosa-LSL-Cas9-eGFP* mouse line, along with AAV-approaches to delete *Lhx2* we found that its loss resulted in decreased morphological complexity (**Fig. 6j-l and Extended Data Fig. 10c,g**). Given its biochemical relationship with NFIA and its role in astrocyte morphogenesis, we determined whether loss of *Lhx2* affects *Gabbr1* expression. To evaluate *Gabbr1* expression we used RNAscope, finding that its expression is significantly reduced in astrocytes that have lost *Lhx2* (**Fig. 6m**). These data illustrate a role for *Lhx2* in promoting astrocyte morphogenesis, and indicates that Lhx2 cooperates with NFIA to regulate *Gabbr1* expression and drive morphogenesis in developing cortical astrocytes.

## Discussion

The cellular and molecular mechanisms by which neuronal input contributes to astrocyte development are fundamental questions. In this study, we demonstrate that astrocyte morphogenesis in the developing cortex is driven by the activity of inhibitory neurons. We further show that deletion of *Gabbr1*, a GABA receptor, in astrocytes results in defective morphogenesis, indicating that it functions as a central regulator of astrocytogenesis. Mechanistically, the link between *Gabbr1* and *Ednrb* reveals new insights into how inhibitory inputs drive signaling pathways that remodel cellular architecture associated with morphology^23, 24^. Endothelin ligands^32^ are released by several cellular sources, further highlighting the role of cell-cell communication as a central driver of astrocyte morphology. Similar to activity-dependent myelination^33–35^, our results indicate that inhibitory neurons provide cues that drive astrocyte development, they also suggest that other forms of activity-dependent input contribute to astrocyte maturation, including excitatory neurons. Given the proximity of peripheral astrocyte processes to neuronal synapses, a model emerges, where astrocyte morphogenesis is likely tuned to the activity of the surrounding neuronal milieu or neurons from a common ancestral origin.

Our finding that *Gabbr1* exhibits region-specific regulation by SOX9 and NFIA, places it as part of the transcriptional program driving astrocytogenesis. Furthermore, we identified new roles for Sox9 and NFIA in astrocyte morphogenesis in the brain, while establishing a new mechanism by which these transcription factors enable developing astrocytes to respond to neuronal cues. Critically, these findings highlight region-specific mechanisms of astrocyte development, where the OB requires SOX9, while the cortex and hippocampus require NFIA. Parallel observations were made in adult astrocytes, indicating that these region-specific transcriptional dependencies in the adult are developmentally encoded^26, 27^. Our studies suggest a mechanism by which transcription factors with region restricted patterns of expression (i.e. LHX2 and NPAS3) confer the regional dependency of ubiquitously expressed transcription factors (i.e. NFIA and SOX9). Together, this suggests a combinatorial transcription factor code, akin to pattern formation, that operates in a region-specific manner to oversee astrocyte development and function.

## Acknowledgements

This work was supported by US National Institutes of Health grants NS071153, AG071687, and NS096096 to BD. We are thankful for support from the David and Eula Wintermann Foundation. We thank Bernhard Bettler for providing the *Gabbr1*-floxed mouse line. scRNA-Seq studies were performed at the Single Cell Genomics Core at BCM partially supported by NIH shared instrument grants (S10OD023469, S10OD025240) and P30EY002520. This project was supported by the Cytometry and Cell Sorting Core at Baylor College of Medicine with funding from the CPRIT Core Facility Support Award (CPRIT-RP180672), the NIH (CA125123 and RR024574) and the assistance of Joel M. Sederstrom. Research reported in this publication was supported by the Eunice Kennedy Shriver National Institute of Child Health & Human Development of the National Institutes of Health under Award Number P50HD103555 for use of the Microscopy Core facilities and the Animal Phenotyping & Preclinical Endpoints Core facilities.

## Authors Contributions

YTC and BD conceived the project and designed the experiments; YTC, JW, ZFL and ELF performed the experiments; JW executed the electrophysiology studies; YTC and ASH designed and executed the bioinformatics analyses. YTC and BD wrote the manuscript.

## Methods

### Animals

All experimental animals were treated in compliance with the US Department of Health and Human Services, the NIH guidelines, and Baylor College of Medicine IACUC guidelines. All mice were housed with food and water available ad libitum in a 12-hour light/dark, 20-22 degree, and 40-60% humidity environment. Both female and male mice were used for all experiments, and littermates of the same sex were randomly allocated to experimental groups. For ex vivo and in vivo experiments, P28 animals were used unless otherwise described. All mice used in this study were maintained on the C57BL/6J background. Different conditional knockout mice were generated by crossing fl/fl mice with Aldh1l1-CreER (The Jackson Laboratory; RRID:IMSR JAX:029655). For *Gabbr1* conditional knockout mice, *Gabbr1^fl/fl^* conditional mutant mice were crossed with Aldh1l1-CreER, resulting in *Gabbr1^fl/fl^*; Aldh1l1-CreER (*Gabbr1 cKO*) and *Gabbr1^fl/fl^* (*Gabbr1 control*) littermate controls^36^. For *Sox9* conditional knockout mice, *Sox9^fl/fl^* conditional mutant mice were crossed with Aldh1l1-CreER, resulting in *Sox9^fl/fl^*; Aldh1l1-CreER (*Sox9 cKO*) and *Sox9^fl/fl^* (*Sox9 control*) littermate controls^37^. For *Nfia* conditional knockout mice, *Nfia^fl/fl^* conditional mutant mice were crossed with Aldh1l1-CreER, resulting in *Nfia^fl/fl^*; Aldh1l1-CreER (*Nfia cKO*) and *Nfia^fl/fl^* (*Nfia control*) littermate controls^26^. For histological analysis, the Aldh1l1-GFP mouse was crossed with control or knockout, resulting in cKO; Aldh1l1-GFP and control; Aldh1l1-GFP mice. For Ca2+ image analysis, Ai96 (RCL-GCaMP6s) mouse (The Jackson Laboratory; RRID:IMSR_JAX:024106) were crossed with control of knockout. For Tdtomato astrocyte labeling, Ai14 (RCL-Td) mouse (The Jackson Laboratory; RRID:IMSR_JAX:007914) were crossed with control of cKO. To induce deletion of *Gabbr1*, *Sox9*, or *Nfia* in developing astrocytes in the P28 brain, P0 pups were injected subcutaneously with 100 mg/kg body weight of Tamoxifen (Sigma-Aldrich, cat no. T5648) dissolved in a 9:1 corn oil/ethanol mixture for single injection at P0-P1. To perform CRISPR-dependent tissue specific knockout of *Ednrb* and *Lhx2*, we utilized Rosa26-LSL-Cas9 knockin mice (The Jackson Laboratory; RRID:IMSR_JAX: 026175). To conditionally knock out *Ednrb* and *Lhx2*, we intraventricularly injected AAV2/9 GFAP-Cre and AAV2/9 U6-sgRNA-GFAP-mCherry into P0-P1 Rosa26-LSL-Cas9 heterozygous and wild-type littermates. Four weeks after injection, we collected the brain, confirmed the expression of EDNRB or LHX2 through immunofluorescence staining, and performed astrocyte morphological analysis and *Gabbr1* RNAscope. For the Imaris analysis, we compared the morphological complexity in two different ways. First, we compared mCherry+ cells in control and mCherry+/Cas9-EGFP+ cells in cKO (shown in **Figure 4f,g**). Second, we compared mCherry−/Cas9-EGFP+ cells and mCherry+/Cas9-EGFP+ cells in cKO (shown in **Extended Data Fig 10d,e**). For EDNRB or LHX2 expression and *Gabbr1* RNAscope, we compared mCherry+ cells in control and mCherry+/Cas9-EGFP+ cells in cKO (shown in **Fig. 6k** and **Extended Data Fig 10b,c**). Above experiments were approved by Baylor College of Medicine IACUC.

### Immunofluorescence on frozen brain tissues

Mice were anesthetized under isoflurane inhalation and perfused transcardially with 1XPBS pH 7.4 followed by 4% paraformaldehyde (PFA). Brains were removed, fixed in 4% PFA overnight, and placed in 20% sucrose for 24 hours before embedded in OCT. Sections of 20 mm (morphological analysis using GFP labeling) were made on a cryostat, washed with 1XPBS 5 min X2, incubated in antigen retrieval buffers at 75 degree 10 min, blocked with 10% goat or donkey serum in PBS with 0.3% Triton x-100, and incubated with primary antibodies in blocking solution overnight. On the next day, sections were incubated with secondary antibodies in PBS with 0.1% Triton x-100 for 1 h RT, followed by incubation with DAPI in PBS for 10min, and mounted with VECTASHIELD Antifade Mounting Media (Vector Laboratories, H-1000). The following primary antibodies were used: Chicken anti-GFP (1:1000; Abcam, ab13970), rabbit anti-NFIA (1:500; Sigma, HPA006111), chicken anti-GFAP (1:1000; Abcam, ab4674), mouse anti-GFAP (1:1000; EMD Millipore, MAB360), goat anti-SOX9 (1:750; RD system, AF3075), rabbit anti-SOX9 (1:650; EMD Millipore, AB5535), rabbit anti-BRN2 (1:1000; Cell Signaling Technology, 12137S), rat anti-CTIP2 (1:500; Abcam, ab18465), rabbit anti-FOXP2 (1:500; Abcam, ab16046), mouse anti-GAD67 (1:200; EMD Millipore, MAB5406), mouse anti-Gephyrin (1:600; Synaptic Systems, 147011), rat anti-HA (1:100; Sigma, 11867423001), rabbit anti-PSD95 (1:200; Thermo Fisher Scientific, 51-6900), guinea pig anti-VGAT (1:350; Synaptic Systems, 131004), guinea pig anti-VGlut1 (1:2000; EMD Millipore, AB5905), guinea pig anti-VGlut2 (1:5000; EMD Millipore, AB2251), rabbit anti-EDNRB (1:250, Abcam, ab117529), rabbit anti-LHX2 (1:250, Abcam, ab184337), rabbit anti-NPAS3 (1:250; Thermo Fisher, PA5-20365). The following secondary antibodies were used (1:500): Alexa Fluor 488 goat anti-chicken (Thermo Fisher Scientific, A11039), Alexa Fluor 568 goat anti-rabbit (Thermo Fisher Scientific, A11036), Alexa Fluor 568 donkey anti-goat (Thermo Fisher Scientific, A11057), Alexa Fluor 568 goat anti-rat (Thermo Fisher Scientific, A11077), Alexa Fluor 488 goat anti-mouse (Thermo Fisher Scientiric, A32723), Alexa Fluor 647 goat anti-guinea pig (Thermo Fisher Scientific, A21450).

### Confocal imaging and image analysis

To evaluate astrocyte morphology, fluorescent images were acquired using a Zeiss LSM 880 laser scanning confocal microscope with 63X oil immersion objective with frame size at 1024 x 1024 and bit depth at 12 (Zen3.1) or a Leica TCS SP8 STED microscope with 63X oil immersion objective with frame size at 1024 x 1024 (LAS X). Serial images at z axis were taken at an optical step of 0.5 mm, with overall z axis range encompassing the whole section. Images were imported to Imaris Bitplane software, and only astrocytes with their soma between the z axis range were chosen for further analysis^38^. We performed 3D surface rendering (**Fig. 1c-e,g-i, 2c,g-i, 4g-i, 6b,j-l, Extended Data Fig. 1c,g, 3a-b,d,e, 8a-d, 10f,g**) using the Imaris Surface module, and color coded the reconstructed surface images based on the surface area of each astrocyte. Morphological analysis was performed using the Imaris Filament module. Astrocyte branches and processes were outlined by Autopath with starting point set at 8 mm and seed point set at 0.7 mm, and statistical outputs including ‘‘filament number Sholl intersections’’ were extracted and plotted with Prism software. Data were generated from 3 brain sections per region per mouse with 3 mice per genotype. The number of astrocytes analyzed were as follows: Gabbr1 control: OB 43, CX 27, HC 32, BS 28, CB 29; Gabbr1 cKO: OB 31, CX 33, HC 30, BS 32, CB 29; Sox9 control: OB 29, CX 35, HC 32, BS 36, CB 32; Sox9 cKO: OB 39, CX 33, HC 24, BS 24, CB 37; Nfia control: OB 56, CX 52, HC 64, BS 28, CB 47; Nfia cKO: OB 60, CX 48, HC 65, BS 43, CB 55; Interneuron Gq Saline: CX 49; Interneuron Gq CNO: CX 49; Interneuron Gi Saline: CX 44; Interneuron Gi CNO: CX 71; CNO only: CX 39; P60 Interneuron Gq Saline: CX 26; P60 Interneuron Gq CNO: 35; Gabbr1 Td control: CX 32, HC 30; Gabbr1 Td cKO: CX 36, HC 38; *sgEdnrb* control: CX 53: *sgEdnrb* cKO: CX 49; *sgLhx2* control: CX 40; *sgLhx2* cKO: CX 42. To analyze number of astrocytes and knockout efficiency of SOX9 and NFIA, fluorescent images were acquired using a Zeiss LSM 880 laser scanning confocal microscope with 20X objective. Cell numbers were quantified by the QuPath software Cell Detection function^39^. To measure the fluorescent intensity of GFAP, fluorescent images were acquired using a Leica TCS SP8 STED microscope with 20X objective or a Zeiss LSM 900 laser scanning confocal microscope with 40X oil objective and were analyzed by Fiji. The person who analyzed the images was blinded to the experimental groups.

### RNAscope

Brain sections were acquired as described above and processed following the sample preparation of fixed frozen tissues of RNAscope® Multiplex Fluorescent Reagent Kit v2 (Advanced Cell Diagnostics, 323100). The mouse *Gabbr1* probe was applied on brain sections (Advanced Cell Diagnostics, 425181). After RNAscope incubation, the sections were then immunostained for astrocyte markers as described above. The images were acquired using a Leica TCS SP8 STED microscope with 63X oil immersion objective with frame size at 1024 x 1024. Serial images at z axis were taken at an optical step of 0.5 mm, with overall z axis range encompassing the whole section. The quantification of *Gabbr1* transcripts number and intensity were analyzed by Fiji.

### EdU cell proliferation assay

P5 pups were Intraperitoneally injected with 100 mg/Kg EdU (Thermo Fisher Scientific, C10337 or C10638) and collected at P7. Brain tissue was processed as described above. After antigen retrieval, sections were washed with 10% goat serum in PBS for 5 minutes and applied Click-iT solution as described in the kit. After 30 minutes EdU staining, sections were washed with 10% goat serum in PBS for 5 minutes, then proceed to immunostaining with desired markers. Images were acquired using Zeiss Axio Imager.M2 with apotome and 20X objective. To quantify proliferating astrocytes, colocalization of EdU and astrocyte markers were analyzed by QuPath software.

### Slice recording

Animals were deeply anesthetized with isoflurane. After decapitation, the brain was quickly excised from the skull and submerged in an ice-cold cutting solution that contained (in mM): 130 NaCl, 24 NaHCO_3_, 1.25 NaH_2_PO_4_, 3.5 KCl, 1.5 CaCl_2_, 1.5 MgCl_2_, and 10 D(+)-glucose, pH 7.4. The whole solution was gassed with 95 % O2-5 % CO_2_. After trimming the hippocampal brain, 300 mm para-sagittal slices were cut using a vibratome with a blade and transferred to extracellular ACSF solution (in mM): 130 NaCl, 24 NaHCO_3_, 1.25 NaH_2_PO_4_, 3.5 KCl, 1.5 CaCl_2_, 1.5 MgCl_2_, and 10 D(+)-glucose, pH 7.4. Slices were incubated at room temperature for at least one hour prior to recording before being transferred to a recording chamber that was continuously perfused with ASCF solution (flow rate = 2 ml/min) Slices were placed in a recording chamber and target cells were identified via upright Olympus microscope with a 60X water immersion objective with infrared differential interference contrast optics. Whole cell recording was performed with pCLAMP10 and MultiClamp 700B amplifier (Axon Instrument, Molecular Devices) at room temperature from layer II-III cortical neurons. The holding potential was −60 mV. Pipette resistance was typically 5-8 MU. The pipette was filled with an internal solution (in mM): 140 K-gluconate, 10 HEPES, 7 NaCl, and 2 MgATP adjusted to pH 7.4 with CsOH for action potential and passive conductance measurements; 135 CsMeSO4, 8 NaCl, 10 HEPES, 0.25 EGTA, 1 Mg-ATP, 0.25 Na_2_-GTP, 30 QX-314, pH adjusted to 7.2 with CsOH (278-285 mOsmol) for EPSC measurement; 135 CsCl, 4 NaCl, 0.5 CaCl_2_, 10 HEPES, 5 EGTA, 2 Mg-ATP, 0.5 Na_2_-GTP, 30 QX-314, pH adjusted to 7.2 with CsOH (278-285 mOsmol) for IPSC measurement. Spontaneous EPSCs were measured in the presence of GABA_A_R antagonist, bicuculline (20 μM, Tocris). IPSCs were measured in the presence of ionotropic glutamate receptor antagonists, APV (50 μM, Tocris), and CNQX (20 μM, Tocris). All holding potential values stated are after correction for the calculated junction potential offset of 14 mV. Electrical signals were digitized and sampled at 50 μs intervals with Digidata 1550B and Multiclamp 700B amplifier (Molecular Devices, CA, USA) using pCLAMP 10.7 software. Data were filtered at 2 kHz. The recorded current was analyzed with ClampFit 10.7 software.

### Two-photon GCaMP imaging in slices

For two-photon imaging, mice were deeply anesthetized with isoflurane and then perfused with cold artificial cerebrospinal fluid (ACSF, in mM:125 NaCl, 25 glucose, 25 NaHCO_3_, 2.5 KCl, 2 CaCl_2_, 1.25 NaH_2_PO_4_ and 1 MgCl_2_, pH 7.3, 310–320 mOsm). The brain was dissected and placed in an ice-cold ACSF. 300 mm thick brain slices were sectioned on a vibratome. Slices were then recovered in oxygenated ACSF (37C) for 15 min and allowed to acclimate to room temperature for at least 15 min before imaging. We recorded calcium traces using a two-photon resonant microscope (LSM 7MP, Zeiss) equipped with a Coherent Chameleon Ultra (II) Ti-sapphire laser tuned to 900 nm and a 20x, 1.0 NA Zeiss objective. Calcium activity was typically sampled at ~1 Hz. Optical signals were recorded for ~5 minutes per trial at 1024 x 1024 pixel resolution. We recorded data from astrocytes at depths of ~30 mm below the surface. All multiphoton imaging experiments were performed within 2-4 hours of slicing. For drug induced calcium imaging, optical signals were recorded after slices were bathed in 500 nM terodotoxin (TTX) for 5 minutes. After 2 minutes of recording under TTX treatment, brain slices were bathed in 50 μM (R)-baclofen (Tocris, 0796) and recorded. Image analysis of Ca^2+^ Spontaneous or drug-induced Ca^2+^ signal was detected in astrocytes expressing GCaMP6s from the olfactory bulb or cortex. The detection of region of interest (ROI) for soma and microdomain for Ca^2+^ imaging was performed in a semi-auto-mated manner using the GECIquant program as described in a previous study^40^. After thresholding from temporally projected stack images with a maximum intensity projection, a polygon selection was manually drawn around the approximate astrocyte territory of interest, and the selection was added to the ImageJ ROI manager. Note that the assignment of territory was approximate and was not used for analysis. The area criterion was 20 mm^2^ to infinity for soma within the GECIquant ROI detection function. Intensity values for each ROI were extracted in ImageJ and converted to dF/F values. For each ROI, basal F was determined during 40 s periods with no fluctuations. Clampfit 10.7 software was used to detect and measure amplitude and frequency values for the somatic and microdomain transients. We counted the response following with these criteria: amplitude (> 0.5 dF/F), pre-trigger time (3 ms), and minimum duration (5 ms).

### Behavioral tests

We subjected 3-month-old male mice to behavioral tests. All the experimental mice were transferred to the testing room at least 30 min prior to the test. All tests were performed with white noise at ± 60 dB in a designated room. The person performing the tests was blinded to the experimental groups.

#### Three-chamber social interaction test

The three-chamber social interaction test was performed in the arena having three chambers, left, middle and right chambers. On the testing day, each animal was first habituated in the chambers with empty wire cages in the left and right chamber for 10 minutes. After habituation, place either LEGO object or partner mouse in the wire cages randomly. Total interaction time with partner mouse was analyzed by ANY-maze software. All the partner mice were habituated to the wire cages in the testing arena for 1 hour per day for 2 days before the day of testing.

#### Open field test

The open field tests were performed using the Versamax system. The Versamax open field chamber is a square arena (40cm 3 40cm 3 30cm, Accuscan Instruments) enclosed by transparent walls. Each mouse was put into the center of the chamber. Locomotor activity was detected automatically by sensor beams at X, Y, and Z directions. Data were recorded in 15 two-minute blocks for 30 min total and were analyzed and exported with Versadat software.

#### Elevated plus maze

The elevated plus maze test were performed on a 1-meter high “+” shaped apparatus with two open arms and two close arms. Mice were put into the center of plus maze and recorded for 10 minutes. The time that mice spent on the open arms or close arms were analyzed by ANYmaze software.

#### Prepulse inhibition

The prepulse inhibition test were performed using SR-LAB-Startle Response System (San Diego Instrument). Mice were put into the cylinder tube in the SR-LAB-Startle Response System chamber and habituated for 5 minutes. After 5 minutes habituation, mice were acclimated to a background white noise of 70 dB for about 5 min prior to the prepulse inhibition test. Each test consisted of 48 trials comprising of 6 blocks of eight trial types each presented in a pseudo random order. Each block had a “No stimulus” trial used to measure baseline mouse response where no sound was presented, a “acoustic startle response” trial comprised of a 40 ms, 120 dB sound burst, a “prepulse only” trials (74, 78 or 82 dB) comprising of three different 20 ms prepulses and finally the “prepulse inhibition (PPI)” trials composed of the presentation of one of the three prepulse sounds, 100 ms prior to the 120 dB startle stimulus. The inter-trial interval ranged from 10 s to 20 s, and the startle response was recorded every 1 ms for 65 ms following the onset of the startle stimulus. Percent PPI of the startle response was calculated as follows: 100 − [(response to acoustic prepulse plus startle stimulus trials/startle response alone trials) × 100].

#### Parallel rod footfall test

The parallel rod foot slip test was performed in a chamber with metal grid floor. For 10 minutes recording, mice freely moved in the chamber. When mice foot slipped on the floor, the ANY-maze software counted as one footfall. The recorded data were analyzed by ANY-maze software.

#### Rotarod

The rotarod test were performed on a rotating rod. It’s a 2-day protocol consisting of 4 trials per day. Each trial lasted for 5 minutes with the rod accelerating at a speed of 4–40 rpm in 5 minutes. The time spent walking on the rod was recorded. Intertrial interval was at least 10–15 minutes.

#### Contextual/cued conditional fear

The contextual conditional fear test was performed in a chamber with metal grid floor. Three checkerboard pattern visual cues (13 cm X 13 cm) were posted at three sides of the chamber. On day 1, mice were put into the center of the chamber and allowed to move freely for 3 min before being exposed to 3 mild foot shocks (2 s, 0.7mA) with 2 min intertrial intervals (ITI) between each shock (figure). On day 2, mice were first put back to the same chamber and movements of mice over 5 min were recorded and analyzed by FreezeFrame software (Actimetrics, Coulbourn Instruments) with the bouts and threshold both set at 6.0 s. % freezing time identified based on the above criteria. Two hours after contextual conditional fear, mice were put back to chamber with different context and were recorded % freezing time upon cue stimulation. The % freezing time in cued conational fear was analyzed by same criteria as contextual conditional fear. Data were then plotted as shown in Figure.

### FACS sorting

We harvested different regions from mouse brains and dissociated them using the protocol described previously^14^. Dissociated astrocytes from different regions were gated with BD FACSDiva Software and sorted by BD FACSAira III with 100 mM nozzle. Around 95,000 GFP^+^ astrocytes were collected per 1.5 mL tube, which contained 650 μl of Buffer RLT (QIAGEN Cat. No. 79216) with 1% b-Mercaptoethanol. Finally, each sample was vortexed and rapidly frozen on dry ice.

### Tissue dissociation for single cell sequencing

Brain slices were prepared as we described in slicing recording methods. The desired brain region was micro-dissected in ACSF on ice, followed by tissue dissociation using neural tissue dissociation kit (Miltenyi Biotec). After 30 minutes incubation on gentleMACS (Miltenyi Biotec), samples were treated with debris removal kit, 1X red blood cell lysis buffer, dead cell removal kit (Miltenyi Biotec) to purify single cells. To remove microglia in samples, CD11b microbeads were applied. Finally, samples were subject to single cell RNA-sequencing library preparation.

### RNA extraction, library preparation and sequencing

For the whole transcriptomic RNA-sequencing, RNA was extracted from pelleted cells using RNeasy Micro Kit (Cat. No. 74004, QIAGEN). RNA integrity (RIN R 8.0) was confirmed using the High Sensitivity RNA Analysis Kit (DNF-472-0500, Agilent formerly AATI) on a 12-Capillary Fragment Analyzer. cDNA synthesis and Illumina sequencing libraries with 8-bp single indices were constructed from 10 ng total RNA using the Trio RNASeq System (0507-96, NuGEN). The resulting libraries were validated using the Standard Sensitivity NGS Fragment Analysis Kit (DNF-473-0500, Agilent formerly AATI) on a 12-Capillary Fragment Analyzer and quantified using Quant-it dsDNA assay kit (Cat. Q33120). Equal concentrations (2 nM) of libraries were pooled and subjected to paired-end (R1: 75, R2: 75) sequencing of approximately 40 million reads per sample using the High Output v2 kit (FC-404-2002, Illumina) on a NextSeq550 following the manufacturer’s instructions. For single-cell RNA-sequencing, single cell gene expression library was prepared according to Chromium Single Cell Gene Expression 3v3.1 kit (10x Genomics). In Brief, single cells, reverse transcription (RT) reagents, Gel Beads containing barcoded oligonucleotides, and oil were loaded on a Chromium controller (10x Genomics) to generate single cell GEMS (Gel Beads-In-Emulsions) where full length cDNA was synthesized and barcoded for each single cell. Subsequently the GEMS are broken and cDNA from each single cell are pooled. Following cleanup using Dynabeads MyOne Silane Beads, cDNA is amplified by PCR. The amplified product is fragmented to optimal size before end-repair, A-tailing, and adaptor ligation. Final library was generated by amplification. Equal concentrations (2 nM) of libraries were pooled and subjected to paired-end (R1: 26, R2: 50) sequencing using the High Output v2 kit (FC-404-2002, Illumina) on a NextSeq550 following the manufacturer’s instructions.

### RNA-Seq bioinformatics analysis

Sequencing files from each flow cell lane were downloaded and the resulting fastq files were merged. Quality control was performed using fastQC (v0.10.1) and MultiQC (v0.9)^41^. Reads were mapped to the mouse genome mm10 assembly using STAR (v2.5.0a)^42^. RNA-seq analysis were analyzed and plotted as previously described^26^. RNA-Seq data can be found at the NIH GEO database (GSE198632).

### Single cell RNA-seq analysis

Sequencing files from each flow cell lane were downloaded and the resulting fastq files were merged. Reads were mapped to the mouse genome mm10 assembly using 10X Cell Ranger (3.0.2) and it is estimated 15,000-38,000 mean reads per cell. For single cell sequencing analysis, standard procedures for filtering, mitochondrial gene removal, doublets removal, variable gene selection, dimensionality reduction, and clustering were performed using Seurat (version 4.1.0) and DoubletFinder^21, 43^. Criteria for cell inclusion were minimum nUMI/cell threshold 200, minimum gene/cell threshold 250, minimum log10gene/UMI threshold 0.8, maximum mitochondria ratio 0.3, and minimum ribosome ration 0.01^44^. Mitochondrial genes were removed before doublets removal. Principle component analysis and elbowplot were used to find neighbors and clusters (resolution 0.3). Cells were visualized using a 2-dimensional Uniform Manifold Approximation and Projection (UMAP) of the PCprojected data. Molecularly distinct cell populations were assigned to each cluster using singleR with adult mouse cortical cell taxonomy single cell RNA-seq data as references^45, 46^. FindAllMarkers were used to identify all differentially expressed markers between clusters. Annotated clusters were refined based on those unique markers. Differentially expressed genes (DEGs) in neurons between Gabbr1 control and cKO were identified by identified by FindMarkers using default settings. 2021 KEGG mouse pathway analysis of DEGs were performed using enrichR^47^. Single cell RNA-Seq data can be found at the NIH GEO database (GSE198357, GSE198633).

### Inference and analysis of cell–cell communication

Cell-cell communications between astrocytes and neurons were inferred using CellChat algorithm^22^. We followed the CellChat workflow, and first identified cell type specific communication within Gabbr1cKO and control experiments separately. Next, we used CellChat to compare the total number of interactions and interaction strength of the inferred cell-cell communication networks in Gabbr1cKO and control experiments. We used netVisual_diffInteraction function to visualize differential number of interactions or interaction strength among Gabbr1cKO and control conditions. Finally, we identified the upgulated and down-regulated signaling ligand-receptor pairs in Gabbr1cKO compared to the control dataset using netVisual_bubble function.

### Co-immunoprecipitation and western Blot

Animal tissues were dissected, washed with cold PBS three times, and dissociated using a pellet homogenizer. RIPA lysis buffer (50 mM Tris-HCl pH 7.5, 150 mM NaCl, 0.5% sodium deoxycholate, 0.1% SDS, 1% NP-40) was used for preparing input control lysates. For co-IP, nuclear lysates were prepared using NE-PER Nuclear and Cytoplasmic Extraction Reagents (ThermoFisher, 78833) according to the manufacturer’s instructions. Brain tissues from 6-8 animals were pulled together to have enough nuclear lysates for each IP. 3mg of nuclear lysates were used for IP with 2 μg IgG or 5 μg primary antibodies (normal mouse IgG, Santa Cruz Biotechnology, sc-2025; normal rabbit IgG, R&D Systems, AB-105-C; mouse anti-LHX2, Santa Cruz Biotechnology, sc-81311; rabbit anti-SOX9, EMDMillipore, ab5535) for overnight at 4 °C. Protein A agarose beads (ThermoFisher, 15918-014) was added for subsequent pull-down for an additional 4 h at 4 °C. The beads were collected, washed, and boiled in 2× SDS gel loading dye to elute immunoprecipitated proteins for Western blot analysis. Input control lysates (20 ug) and immunoprecipitated proteins were run on a 8% SDS polyacrylamide gel, followed by transferring onto nitrocellulose membrane at 350 mA for 65 min. 5% milk in Tris-buffered saline with Tween20 (TBST) was used to block the membrane, followed by incubating primary antibodies at 1:1,000 dilution (rabbit anti-LHX2, Abcam, ab184377; rabbit anti-NFIA, Sigma, HPA006111; rabbit anti-SOX9, EMDMillipore, ab5535; rabbit anti-NPAS3, ThermoFisher, PA520365) for overnight at 4 °C. The next day, membranes were washed three times with TBST, incubated with horseradish peroxidase-conjugated anti-rabbit or anti-mouse IgG at 1:2,000 dilution in 5% milk in TBST at room temperature for 1 h. Membranes were then washed with TBST three times before developing with luminol reagent (Santa Cruz Biotechnology, sc-2048).

### Chromatin immunoprecipitation PCR (ChIP-PCR)

Mouse cortex and olfactory bulbs were collected for ChIP experiments. Dissociated cortexes or olfactory bulbs were pooled together from 2-3 animals for each ChIP experiment. Chromatin was crosslinked by using freshly prepared 1.1% formaldehyde solution with rocking at room temperature for 10 min, followed by addition of 0.1 M glycine. Cell pellets were collected by centrifugation at 3500 rpm for 5 min at 4 C, washed with PBS and frozen at 80 C or used immediately for preparing lysates. Pellets were resuspended with PBS/PMSF containing 0.5% Igepal to release nuclei, followed by washing with cold ChIP-Buffer (0.25% TritonX, 10 mM EDTA, 0.5 mM EGTA, 10 mM HEPES pH 6.5) and nuclei were lysed with ChIP lysis buffer (0.5% SDS, 5 mM EDTA, 25 mM Tris-HCl pH 8) for 15-20 min at room temperature. Lysates were sonicated to 250-350 bp using Diagenode Bioruptor. Immunoprecipitation was carried out by rotating sonicated lysates overnight at 4 C with NFIA antibody (5 mg, Sigma, HPA006111) or SOX9 antibody (EMD Millipore, AB5535) followed by pull-down using Protein A/G agarose beads (Thermo Fisher Scientific, 15918014) for 6 hours. The beads were collected and washed with TSE1 buffer (0.1% SDS, 1% TritonX, 2 mM EDTA, 20 mM Tris-HCl pH 8, 150 mM NaCl), TSE2 buffer (TSE1 buffer with 500 mM NaCl), LiCl buffer (0.25M lithium chloride, 1% NP40, 1% sodium deoxycholate, 1 mM EDTA, 10 mM Tris-HCl pH 8) and TE buffer. Immunoprecipitated chromatin was then eluted by heating the beads in ChIP Elution buffer (1% SDS, 0.1 M NaHCO3) at 65 C for 20 min twice. A small sample of elution was used for Western Blot analysis to confirm immunoprecipitation of NFIA. ChIP-DNA was quantified using Quant-it dsDNA assay kit (Cat. Q33120) and used for ChIP-PCR. Primers for NFIA binding motif and SOX9 on Gabbr1 promoter: Forward 5’-TTCAAGGTCTGTTCCCCAGGC-3’, Reverse 5’-GAGGGCGTAGAGGTAGGATGGA-3’.

### Intraventricular injection of AAV viruses

For interneuron hM3Dq experiments, we used AAV2/9-pAAV-hDlx-hM3Dq-dTomato-Fishell-4 (Addgene, 83897) and AAV2/9-pAAV-hDlx-hM4Di-dTomato-Fishell-5 (Addgene, 83896) at a concentration of 6+13 genome copies per ml (gc/ml). For the GCaMP experiments in *Sox9* olfactory bulb experiments, we generated pAAV-GFAP-GCaMP6m plasmid from flexed-GCaMP6 and pZac2.1-GfaABC1D-mCherry−hPMCA2w/b (Addgene, 111568) and used AAV2/9-pAAV-GFAPGCaMP6m at a concentration of 1.07527E+13 genome copies per ml (gc/ml). For interneuron labeling experiments, we generated pAAV-mDlxRuby2 plasmid from pAAV-mDlx-NLS-Ruby2 (Addgene, 99130) and used AAV2/9-pAAV-mDlx-Ruby2 at a concentration of 2.344E+13 genome copies per ml (gc/ml)^48^. For intraventricular injection, P1 pups were anesthetized with hypothermia and injected AAV virus as described in previous paper^49^. 2.5 μl of Trypan Blue (Thermo Fisher Scientific, 15250061) was mixed with 10 μl of virus before injection. AAV virus: All AAV viruses were generated by the Optogenetics and Viral Vectors Core at Jan and Dan Duncan Neurological Research Institute (NRI). For astrocyte, CRISPR-dependent tissue specific knockout experiments, we utilized pZac2.1-U6-sgRNA empty-GfaABC1D-mcherry (Khakh lab) to generate pAAV-U6-Ednrb sgRNA-Gfap-mcherry and pAAV-U6-Lhx2 sgRNA-Gfap-mcherry by amplifying sgRNA inserts with forward primers: CACCGTCAATATTTCGTTGGCACGG (*Ednrb*), CACCGGCTGCACAGAGAACCGCCTG (*Lhx2*) and reverser primers: AAACCCGTGCCAACGAAATATTGAC (*Ednrb*), AAACCAGGCGGTTCTCTGTGCAGCC (*Lhx2*). We used AAV2/9-pAAV-Gfap-Cre-P2A-TurboRFP at a concentration of 5E+12 genome copies per ml (gc/ml) and AAV2/9-pAAV-U6-Ednrb sgRNA-Gfap-mcherry or AAV2/9-pAAV-U6-Lhx2 sgRNA-Gfap-mcherry at a concentration of 2E+12 genome copies per ml (gc/ml). All animal procedures were done in accordance with approved BCM IACUC protocols.

## QUANTIFICATION AND STATISTICAL ANALYSIS

Sample sizes and statistical tests can be found in accompanying Figure legends. Offline analysis was carried out using Clampfit 10.7, Minianalysis, SigmaPlot 13, Prism 9, and Excel software. we do not assume that our data are normally distributed and perform either linear or generalized mixed-effects model for repeated measurements. If the number of paired animals is more than 3, we used Mann-Whitney test based on the number of animals. For the number of paired animal equals to 3, we used either linear or generalized linear mixed-effects model (LME or GLME) to consider the variances from both cells and animals. We chose between LME or GLME based on the data distribution. We used Shapiro-Wilk test to test whether the analyzed cells are normally distributed. If the data is normally distributed, we used LME to perform statistical analysis. If the data is not normally distributed, we used GLME to analyze the data. Data are presented as mean ± SEM (standard error of the mean). Levels of statistical significance are indicated as follows: * (p < 0.05), ** (p < 0.01), *** (p < 0.001), **** (p < 0.0001).

### Reporting summary

Further information on research design is available in the Nature Research Reporting Summary linked to this paper.

### Data availability

The bulk RNA-seq data from developing astrocytes and Ednrb1-cKO astrocytes has been deposited in the NCBI Gene Expression Omnibus (GEO) under accession number GSE198632. Single cell RNA-Seq data can be found at the NIH GEO database (GSE198357, GSE198633). All other data in this article are available from the corresponding author upon reasonable request.

### Code availability

No custom code was used. R package limma eBayes function was used to define differentially expressed genes. Bioconductor SVA/Combat package was used for batch correction.

## Extended Data Figure Legends

**<u>Extended Data Figure 1. Normalization of astrocyte morphology in the adult after developmental activation of inhibitory neurons</u>**

**a-b.** Analysis of SOX9 (e) and Ki67(f) expression within Aldh1l1-GFP astrocytes at P21 after activation of inhibitory neurons (or control); quantification was derived from *n* = 3 pairs of animals (**a**, 20,24 images; GLME; **b**, 12, 11 images; GLME). **c.** CNO only treatment of Aldh1l1-GFP mice from P7-P21 and analysis of astrocyte morphology at P21. *n* = 3 animals (39 cells; GLME model with Sidak’s multiple comparisons test). **d.** Heatmap depicting expression of GABA receptor subunits in developing astrocytes from the cortex (CX), hippocampus (HC), or olfactory bulb (OB) at P1, P7, and P14. See Extended Data Figure 2. **d.** Example of gating strategy and percentage of GFP^+^ astrocytes FACS isolated from P1 animal. **e.** Heatmap depicting expression of GABA receptor subunits in developing astrocytes from the cortex (CX), hippocampus (HC), or olfactory bulb (OB) at P1, P7, and P14. See Extended Data Figure 2. **f.** Schematic of DREADD-based activation of inhibitory neurons in post-natal Aldh1l1-GFP mice and experimental timeline. **g.** Imaging of P60 Aldh1l1-GFP astrocytes after hM3Dq activation of inhibitory neurons; quantification of morphological complexity using Scholl analysis, branch number, and total processes at P21; *n* = 3 pairs of animals (26,35 cells; upper, GLME model with Sidak’s multiple comparisons test; bottom, GLME model). Scale bars, 20 μm (**a-c**), 30 μm (**g**). Data represent mean ± s.d. (**a-c**, **g** upper), median, minimum value, maximum value and IQR (**g** bottom).

**<u>Extended Data Figure 2. Transcriptomic RNA-Sequencing analysis of developing astrocytes in the cortex, hippocampus, and olfactory bulb at P1, P7, and P14.</u>**

**a.** Heatmap depicting the expression of neuron-specific and astrocyte-specific genes from P1, P7, and P14 FACS isolated Aldh1l1-GFP astrocytes from the listed brain regions. **b.** Aldh1l1-GFP astrocytes from the cortex at P1, P7, P14. Principal component (PC) analysis against top 2,000 variable genes across the region and timepoints examined from 3-4 animals in each group. **c.** Heatmap depicting differential patterns of gene expression in developing astrocytes across brain regions and timepoints. **d**. Gene Ontology (GO) analysis of the common and region-specific patterns of gene expression. **e-f.** *Ald1l1-CreER; ROSA-LSL-tdTomato* mouse line treated with tamoxifen at P1, harvested at P28. Co-immunostaining of tdTomato labeled cells with Sox9, Olig2, NeuN, and Iba1 demonstrating astrocyte-specific activity of Aldh1l1-CreER line. *n* = 4 animals. Scale bars, 10 μm (**b**), 40 μm (**e-h**).

**<u>Extended Data Figure 3. Analysis of astrocyte development in the *Gabbr1-cKO* mouse line.</u>**

**a.** Imaging of Aldh1l1-GFP astrocytes from the brain stem and cerebellum at P28; quantification of morphological complexity was derived from *n* = 3 pairs of animals (*Gabbr1 control*: BS 28, CB 29; *Gabbr1 cKO*: BS 32, CB 29 cells; GLME model with Sidak’s multiple comparisons test, \**P* = 0.0179, 0.0167). **b.** Quantitative analysis of branch points and process length from all brain regions analyzed; *n* = 25-38 cells from 3 pairs of animals (*Gabbr1 control*: OB 38, CX 30, HC 29, BS 28, CB 25; *Gabbr1 cKO*: OB 33, CX 30, HC 29, BS 32, CB 29 cells; two way ANOVA, \*\**P* = 0.0014, \**P* = 0.0174, *P* = 0.9040, \**P* = 0.0132, *P* = 0.7126, \*\**P* = 0.0054, \*\*\*\**P* < 0.0001, \*\**P* = 0.0066, *P* = 0.3763). **c.** Schematic describing the experimental timeline and mouse lines rendering astrocyte-specific knockout of *Gabbr1* for sparse labeling experiments. **d-e**. Imaging and quantification of sparsely labeled, tdTomato-expressing astrocytes from *Gabbr1-cKO* and *control* mice from the cortex (d) and hippocampus (e); *n* = 3 pairs of animals (*Gabbr1 control*: CX 32, HC 30; *Gabbr1 cKO*: CX 36, HC 38 cells; **d,e** upper, GLME model with Sidak’s multiple comparisons test, \**P* = 0.0213, \*\**P* = 0.0012; **d,e** bottom, GLME model, \*\*\**P* = 0.00043, \*\**P* = 0.0027, \*\*\**P* = 0.00042, \*\*\*\**P*<0.0001). **f.** Antibody staining for SOX9 in Aldh1l1-GFP astrocytes from cortex of *Gabbr1-cKO* and control; quantification is derived from *n* = 3 pairs of animals (35 images; GLME model). **g.** Pulse-chase EdU-labeling and antibody staining at P28 from all brain regions analyzed; quantification is derived from *n* = 3 pairs of animals (*Gabbr1 control*: OB 9, CX 9, HC 9, BS 9, CB 9; *Gabbr1 cKO*: OB 8, CX 9, HC 9, BS 9, CB 9 images; GLME model). Scale bars, 30 μm (**a**), 20 μm (**d-g**). Data represent mean ± s.d. (**a-b**, **d-e** upper, **f-g**), median, minimum value, maximum value and IQR (**d-e** bottom).

**<u>Extended Data Figure 4. RNA-Seq of *Gabbr1-cKO* astrocytes and single cell RNA-Seq analysis of *Gabbr1-cKO* cortex.</u>**

**a.** Serut analysis of single cell RNA-Seq (scRNA-Seq) from *Gabbr1-cKO* and controls from P28 cortex. **b.** Quantification of cell clusters identifying CNS cell types from scRNA-Seq data. **c-e**. Dot plot summaries demonstrating CellChat interaction profiles and expression patterns of key astrocyte-neuron interaction pathways.

**<u>Extended Data Figure 5. Analysis of cortical neurons in the *Gabbr1-cKO* mouse line and behavioral studies.</u>**

**a.** Antibody staining for BRN2 (Layers II/II). **b.** CTIP2 (Layers V). **c.** FOXP2 (Layers VI) layer-specific neuronal markers in the P28 cortex from *Gabbr1-cKO* and *control*; quantification is derived from *n* = 11-12 images from 3 pairs of animals (*control* 12, *cKO* 11 images; GLME model). **d.** Schematic of synaptic markers and cortical layers. **e-f.** Antibody staining for makers of excitatory synapses Vglut1/PSD95 (e) and Vglut2/PSD95 (f) in layer I of the cortex from *Gabbr1-cKO* or *control* mice at P28 (*n* =3 pairs of animals; GLME model, \**P* = 0.0490). **g.** Antibody staining for markers of inhibitory synapses VGAT/Gephyrin at P28; quantification is derived from 3 pairs of animals (GLME model). **h-m**. Summary of behavioral assays conducted on Gabbr1-cKO and control animals including open field (h), elevated plus maze (i), rotarod (j), parallel foot fall (k), contextual fear conditioning (l), and cued fear conditioning (m).; data is derived from 10 *control* and 11 *Gabbr1-cKO* animals (two-tailed Mann-Whitney test, \*\*\**P* = 0.0004). Scale bars, 100 μm (**a-c**), 3 μm (**e-g**). Data represent mean ± s.d. (**a-c**, **e-m**).

**<u>Extended Data Figure 6. Electrophysiological recordings from cortical excitatory neurons from *Gabbr1-cKO*</u>**

**a.** Two-photon, slice imaging of GCaMP6s activity in *control* and *Gabbr1-cKO* astrocytes from the cortex at P28. Quantification of Ca^2+^ activity in astrocytic microdomains in the *Gabbr1-cKO* and *control* animals, quantification is derived from *n* = 3 pairs of animals (19, 30 cells; GLME model). **b-e.** Representative traces of action potential in layer II/III excitatory neurons upon varying injected current in *Gabbr1-cKO* and *control* (b). Summary data of action potential firing (c; two way ANOVA). Summary data of resting membrane potential (d; two-tailed unpaired Welch’s t-test) and threshold (e; two-tailed unpaired Welch’s t-test) from 3 pairs of animals (*n* = 13, 12 cells). **f-i.** Representative traces of action potential in layer II/III inhibitory neurons upon varying injected current in *Gabbr1-cKO* and *control* (f). Summary data of action potential firing (g; two way ANOWA). Summary data of resting membrane potential (h; two-tailed unpaired Welch’s t-test, \*\**P* = 0.0091) and threshold (i; two-tailed unpaired Welch’s t-test) from 3 pairs of animals (*n* = 12, 15 cells). Scale bars, 20 μm (**a**). Data represent median, minimum value, maximum value and IQR (**a**), mean ± s.e.m. (**c-e**, **g-i**).

**<u>Extended Data Figure 7. Analysis of astrocyte development in the *Sox9-cKO* and *Nfia-cKO* mouse lines.</u>**

**a.** Analysis of NFIA expression in Aldh1l1-GFP astrocytes from the *Nfia-cKO* and *control* at P7 in the cortex, hippocampus, and olfactory bulb; quantification of knockout efficiency was derived from 3 pairs of animals (two-way ANOVA, \*\*\*\**P*<0.0001). **b.** Analysis of SOX9 expression in Aldh1l1-GFP astrocytes from the *Sox9-cKO* and *control* at P7 in the cortex, hippocampus, and olfactory bulb; quantification of knockout efficiency was derived from 3 pairs of animals (two-way ANOVA, \*\*\*\**P*<0.0001). **c.** Analysis of NFIA expression in Aldh1l1-GFP astrocytes from the *Nfia-cKO* and *control* at P28 in the cortex, hippocampus, cerebellum, and olfactory bulb; quantification of knockout efficiency was derived from 3 pairs of animals (two-way ANOVA, \*\*\*\**P*<0.0001). **d.** Analysis of SOX9 expression in Aldh1l1-GFP astrocytes from the *Sox9-cKO* and *control* at P28 in the cortex, hippocampus, cerebellum, and olfactory bulb; quantification of knockout efficiency was derived from 3 pairs of animals (two-way ANOVA, \*\*\*\**P*<0.0001, \**P* = 0.0205). **e-f.** Two-photon, slice imaging of spontaneous GCaMP6s activity in *control* and *Nfia-cKO* astrocytes from the cortex at P28 (e) or *control* and *Sox9-cKO* astrocytes from the olfactory bulb at P28 (f); quantification is derived from 3 pairs of animals (two-tailed Mann-Whitney test). **g**. AAV-based overexpression of NFIA in the developing cortex, analysis of *Gabbr1* expression at P28 via RNAscope; *n* = 3 pairs of animals (19, 18 cells; LME model, \**P* = 0.023, \*\*\**P* = 0.00034). **h**. AAV-based overexpression of SOX9 in the developing olfactory bulb, analysis of *Gabbr1* expression at P28 via RNAscope; *n* = 3 pairs of animals (20,25 cells). Scale bars, 50 μm (**a-d**), 10 μm (**e-f**), 20 μm (**g-h**). Data represent mean ± s.d. (**a-d**), mean ± s.e.m. (**e-f**), median, minimum value, maximum value and IQR (**g**-**h**).

**<u>Extended Data Figure 8. Analysis of astrocyte morphogenesis in the *Sox9-cKO* and *Nfia-cKO* mouse lines.</u>**

**a-b.** Imaging of Aldh1l1-GFP astrocytes from the hippocampus, brainstem, and cerebellum at P28 from the *Nfia-cKO* (a) or *Sox9-cKO* (b) and associated controls; quantification of morphological complexity via Scholl analysis was derived from *n* = 3 pairs of animals (**a**, *Nfia control*: HC 64, BS 28, CB 47; *Nfia cKO*: HC 65, BS 43, CB 55 cells; GLME model with Sidak’s multiple comparisons test, \*\**P* = 0.0015; **b**, *Sox9 control*: HC 32, BS 36, CB 32; *Sox9 cKO*: HC 24, BS 24, CB 37; GLME model with Sidak’s multiple comparisons test). **c-d.** Quantification of astrocytic branch number and process length from *Nfia-cKO* (c) or *Sox9-cKO* (d) across cortex, olfactory bulb, hippocampus, brainstem, and cerebellum; derived from *n* = 3 pairs of animals (**c**, *Nfia control*: OB 59, CX 43, HC 54, BS 27, CB 48; *Nfia cKO*: OB 50, CX 56, HC 59, BS 38, CB 54 cells; two-way ANOVA, \*\**P* = 0.0054, \*\*\*\**P* <0.0001; **d**, *Sox9 control*: OB 29, CX 31, HC 32, BS 37, CB 33; *Sox9 cKO*: OB 40, CX 27, HC 33, BS 21, CB 39; two-way ANOVA, \*\**P* = 0.0025, \*\*\*\**P* < 0.0001, \*\*\**P* = 0.0002). **e-f.** Pulse-chase EdU-labeling and antibody staining at P28 from the cortex of *Nfia-cKO* (e) and olfactory bulb of Sox9-cKO (f); quantification is derived from 3 pairs of animals (two-tailed Mann-Whitney test). **g-h.** Quantification of the number of Aldh1l1-GFP astrocytes in the cortex of the *Nfia-cKO* (g) or olfactory bulb from *Sox9-cKO* and associated controls; quantification is derived from 3 pairs of animals (two-tailed Mann-Whitney test). Scale bars, 30 μm (**a-b**), 50 μm (**e-h**). Data represent mean ± s.d. (**a-h**).

**<u>Extended Data Figure 9. Electrophysiological and behavioral analysis of the *Nfia-cKO* mouse line.</u>**

**a.** Schematic of synaptic markers and cortical layers. **b-c.** Antibody staining for makers of excitatory synapses Vglut1/PSD95 (b) and Vglut2/PSD95 (c) in layer I of the cortex from NFIA-cKO or control mice at P28, quantification is derived from 3 pairs of animals (GLME model). **d.** Antibody staining for markers of inhibitory synapses VGAT/Gephyrin at P28; quantification is derived from 3 pairs of animals (GLME model). **e-f.** Representative traces of spontaneous EPSCs and IPSCs from excitatory (e) and inhibitory (f) neurons from cortex of *Nfia-cKO* and *controls*. Associated cumulative and bar plots demonstrate quantification of amplitude and frequency of sEPSC and sIPSC from 3 pairs of animals (**e**, Kolmogorov-Smirnov test, **** *P* < 0.0001, \**P* = 0.0181, two-tailed Mann-Whitney test; f, Kolmogorov-Smirnov test, \*\**P* = 0.0026, **** *P* < 0.0001, two-tailed Mann-Whitney test, \**P* = 0.0149). **g-h**. 3-chamber social interaction and prepulse inhibition behavioral studies on *Nfia-cKO* and *control* mice from 11-13 animals in each group (**g**, *n* = 13 pairs of animals, two-way ANOVA, \*\*\**P* = 0.0002; **h**, *n* = 13, 11 animals, two-tailed Mann-Whitney test, \*\**P* = 0.0050). **i-j**. Representative traces of action potential in layer II/III neurons upon varying injected current in *Nfia-cKO* and *control*. Summary data of action potential firing, resting membrane potential, and threshold from excitatory neurons (i) and inhibitory neurons (j); quantification is derived from at least 3 pairs of animals (two-way ANOVA and two-tailed Mann-Whitney test). **k-p.** Summary of behavioral tests from NFIA-cKO and control, including open field; *n* = 13, 11 animals (k), elevated plus maze; *n* = 10 animals (l), rotarod; *n* = 5, 8 animals (m), parallel footfall; *n* = 13, 11 animals (n), contextual fear conditioning; *n* = 12, 10 animals (o), and cued fear conditioning; *n* = 12, 10 animals (p) (two-tailed Mann-Whitney test, *P = 0.0265, 0.0136). Scale bars, 3 μm (**b-d**). Data represent mean ± s.d. (**b-d**, **g**, **o-p**), mean ± s.e.m. (**e-f**, **h-n**)

**<u>Extended Data Figure 10. Analysis of cortical neurons in the *Nfia-cKO* mouse line.</u>**

**a.** Antibody staining for BRN2 (Layers II/II), CTIP2 (Layers V), and FOXP2 (Layers VI) layer-specific neuronal markers in the P7 cortex from *Nfia-cKO* and control; quantification is derived from *n* = 3 pairs of animals (6 images, GLME model). **b.** Quantification of EDNRB expression in virally labeled astrocytes from the P28 cortex of mice where it has been knocked out using guideRNAs in the *ROSA-LSL-Cas9-EGFP* mouse line; quantification is derived from *n* = 3 pairs of animals (37, 38 cells; LME model, \*\**P* = 0.0068). **c**. Quantification of LHX2 expression in virally labeled astrocytes from the P28 cortex of mice where it has been knocked out using guideRNAs in the *ROSA-LSL-Cas9-EGFP* mouse line; quantification is derived from *n* = 3 pairs of animals (37, 39 cells; GLME model, \*\*\**P* = 0.00099). **d-e.** Western blots before cropped, arrow heads label the proteins of interest. **f-g.** Imaging of virally labeled astrocytes from the P28 cortex of *Ednrb-cKO* or *Lhx2-cKO* mice where *Ednrb* or *Lhx2* has been knocked out using guideRNAs in the ROSA-LSL-Cas9-EGFP mouse line; quantification of morphological complexity via Scholl analysis was derived from *n* = 3 animals (**f**, *Ednrb-cKO*: 22 mcherry^−^Cas9-EGFP^+^ cells, 49 mcherry^+^Cas9-EGFP^+^ cells; GLME model with Sidak’s multiple comparisons test, \**P* = 0.0307; GLME model, *P* = 0.06, \*\**P* = 0.008; **g**, *Lhx2-cKO*: 13 mcherry^−^Cas9-EGFP^+^ cells, 40 mcherry^+^Cas9-EGFP^+^ cells; GLME model with Sidak’s multiple comparisons test, \*\**P* = 0.0037; GLME model, \*\**P* = 0.006, 0.003). Scale bars, 100 μm (**a**). Data represent mean ± s.d. (**a**, **f-g** left), median, minimum value, maximum value and IQR (**b-c**, **f-g** right).

